# Exploring Ethylene-Related Genes in *Cannabis sativa*: Implications for Sexual Plasticity

**DOI:** 10.1101/2023.04.28.538750

**Authors:** Adrian S. Monthony, Maxime de Ronne, Davoud Torkamaneh

**Affiliations:** Département de phytologie, Université Laval, Québec City, Québec, Canada; Institut de Biologie Intégrative et des Systèmes (IBIS), Université Laval, Québec, Canada; Centre de recherche et d’innovation sur les végétaux (CRIV), Université Laval, Québec, Canada; Institut intelligence et données (IID), Université Laval, Québec, Canada

**Keywords:** ethylene, sexual plasticity, cannabis, STS, biosynthesis, sex expression, sex determination, transcriptomics

## Abstract

Sexual plasticity is a phenomenon wherein organisms possess the ability to alter their phenotypic sex in response to environmental and physiological stimuli, without modifying their sex chromosomes. *Cannabis sativa* L., a medically valuable plant species, exhibits sexual plasticity when subjected to specific chemicals that influence ethylene biosynthesis and signaling. Nevertheless, the precise contribution of ethylene-related genes (ERGs) to sexual plasticity in cannabis remains unexplored. The current study employed *Arabidopsis thaliana* L. as a model organism to conduct gene orthology analysis and reconstruct the Yang Cycle, ethylene biosynthesis, and ethylene signaling pathways in *C. sativa*. Additionally, two transcriptomic datasets comprising male, female, and chemically induced male flowers were examined to identify expression patterns in ERGs associated with sexual determination and sexual plasticity. These ERGs involved in sexual plasticity were categorized into two distinct expression patterns: floral organ concordant (FOC) and unique (uERG). Furthermore, a third expression pattern, termed karyotype concordant (KC) expression, was proposed, which plays a role in sex determination. The study revealed that CsERGs associated with sexual plasticity are dispersed throughout the genome and are not limited to the sex chromosomes, indicating a widespread regulation of sexual plasticity in *C. sativa*.

**Key Message:** Presented here are model Yang cycle, ethylene biosynthesis and signaling pathways in *Cannabis sativa*. *C. sativa* floral transcriptomes were used to predict putative ethylene-related genes involved in sexual plasticity in the species.

## Introduction

*Cannabis sativa* L. (cannabis) belongs to the estimated 5-6% of dioecious angiosperm species (Renner, 2014). Despite their rarity, dioecious plants provide an excellent opportunity to study the mechanisms involved in sex expression and the evolution of sex-determining regions and sex chromosomes. *C. sativa* (2n=20) is predominantly dioecious with a XY sex-determination system; however, monoecious genotypes of *C. sativa* have been bred, especially for the production of seeds and oil used in industrial hemp (Ferfuia *et al*., 2021). Additionally, dioecious cannabis genotypes have been known to display hermaphroditism (Monthony *et al*., 2021c), reversion from a sexual to vegetative state (Monthony *et al*., 2021a), and sometimes complete changes in sex expression in response to certain environmental factors and chemical reagents (Schaffner, 1923; Heslop-Harrison, 1956; Ram & Jaiswal, 1972; Mohan Ram & Sett, 1982; Lubell & Brand, 2018; Moon *et al*., 2020a). Taken together, these observations suggest that cannabis sex is plastic and not strictly controlled by its XX or XY chromosomal complement.

Sexual plasticity has been described as the ability of an organism to change its phenotypic sex in response to environmental or physiological cues without altering its biological sex (Liu *et al*., 2017). Vertebrates exhibit sexual plasticity on a continuum from genotypic sex determination (GSD) to environmental sex determination (ESD), with some organisms influenced by both (GSD + ESD; Lambert *et al*., 2019). Non-mammalian vertebrates, such as fish (Senthilkumaran, 2015; Liu *et al*., 2017) and frogs (Lambert *et al*., 2019), as well as insects like aphids and bees (Morgan, 1909; Charlesworth & Mank, 2010), exhibit sexual plasticity triggered by diverse factors. The mechanisms responsible for these changes vary, but can involve gene expression changes related to hormone synthesis and signaling, germ cell development and maintenance as well as microRNAs (Perry & Grober, 2003; Hayes *et al*., 2010; Ashby *et al*., 2016; Cardoso-Júnior *et al*., 2017). For instance, social factors such as population density, mating competition, and hierarchical structures can affect sexual plasticity in fish (Warner & Swearer, 1991; Liu *et al*., 2017). Amphibians can exhibit sexual plasticity under normal and extreme conditions (Lambert *et al*., 2015, 2019), while in some insects, the mode of sexual reproduction can affect offspring sex (Morgan, 1909; Charlesworth & Mank, 2010). Although the term sexual plasticity has largely been used to describe fluid sex expression in the animal kingdom (Schlesinger *et al*., 2010; Senthilkumaran, 2015; Liu *et al*., 2017), the present study highlights how sexual plasticity can also be used to describe the interaction between GSD and ESD in dioecious plants with sex chromosomes, such as *C. sativa*.

The concept of sexual plasticity, as addressed in this study, describes a departure from the expected sexual traits regulated by the sex chromosomes in dioecious plants. This deviation occurs due to the influence of specific genes, termed sexual plasticity genes, which, when affected by external factors -including chemicals, stresses, changes in photoperiod or climate fluctuations-prompt a shift from the plant’s chromosomal sex to a contrasting phenotypic sex, akin to observed phenomena in vertebrates (Schaffner, 1923; Liu *et al*., 2017; Lambert *et al*., 2019). To date, the control of sexual plasticity in dioecious plants has not been widely described. In contrast, sex determination in many monoecious plants is more widely studied and understood. For example, in Cucurbitaceae, a family of predominantly monoecious plants lacking sex chromosomes, specific loci govern sex determination, where particular alleles or allele combinations directly influence flower sex (Yamasaki *et al*., 2001; Liu *et al*., 2008). This distinction between monoecious and dioecious plants with sex chromosomes has led us to hypothesize that sexual plasticity genes confer adaptive and reproductive flexibility. This adaptation allows plants to manifest variations in sexual expression, permitting them to dynamically adjust their reproductive strategies in response to external cues.

As early as the 1920s, researchers had noted the plasticity of cannabis sex, linking it to environmental factors such as photoperiod (Schaffner, 1923) . However, significant understanding of the factors that induce sexual plasticity stem from informal research by enthusiasts following cannabis criminalization until the recent progression toward legalization. The outcomes of this informal research revealed that chemical exposure can induce sexual plasticity in plants, akin to the plasticity observed in amphibians (Green, 2005; Thomas, 2012). This important early research identified silver-containing compounds such as silver thiosulfate (STS), a potent inhibitor of ethylene signaling, as promoting the production of male flowers on female plants (induced male flowers; IMF) and aqueous ethephon ((2-Chloroethyl) phosphonic acid), which rapidly decomposes to produce ethylene gas, as promoting the production of female flowers on male plants (induced female flowers; IFF) as shown in Figure 1. Following the legalization of cannabis in certain jurisdictions, several research groups (Lubell & Brand, 2018; Kurtz *et al*., 2020; Moon *et al*., 2020a; Flajšman *et al*., 2021) have reconfirmed and quantified the inverse effects of STS and ethephon on cannabis sexual expression. These studies point to ethylene as a key effector of sexual plasticity. Although the phenotypic outcomes of these treatments in cannabis have been quantified, to date there are no studies that have sought to understand the underlying molecular mechanisms controlling ethylene-related sexual plasticity in cannabis.

**Figure 1.**
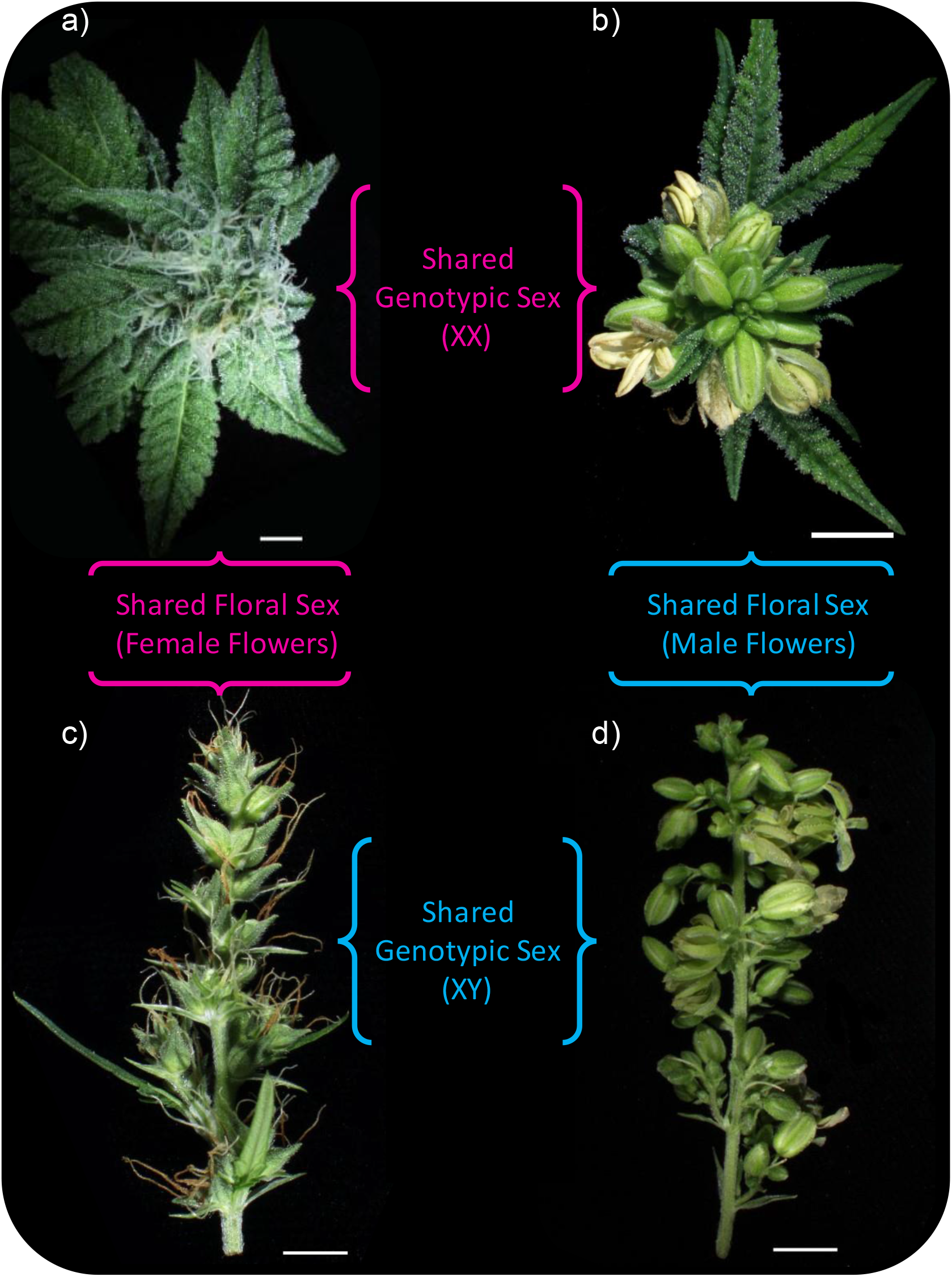
Representative photos of four floral phenotypes in *Cannabis sativa* (cv. La Rosca), grown at Université Laval. a) 4-week-old female flowers on a genetically female plant; b) 4-week-old STS-induced male flowers (IMFs) on a genetically female plant; c) ethephon-induced female flowers (IFFs) on a genetically male plant and d) 3-week-old male flowers on a genetically male plant. Scale bars = 5 mm.

Ethylene was discovered in 1901 and has been extensively studied since then (Coulter & Barnes, 1901; Gane, 1934; Kende, 1998; Chang, 2016). It plays a crucial role as a plant growth regulator in numerous plant processes such as seed germination, root and shoot growth, flowering, fruit ripening, leaf and fruit abscission, as well as responses to environmental stresses and senescence (Tieman *et al*., 2000; Resnick *et al*., 2006; Argueso *et al*., 2007; Yang *et al*., 2015; Dhakal *et al*., 2019; Binder, 2020). Ethylene is synthesized from S-adenosyl methionine (SAM) through a two-step pathway. The first step involves the enzyme 1-aminocyclopropane-1-carboxilic acid (ACC) synthase (ACS), which converts SAM to ACC and 5’-methylthioadenosine (MTA; Argueso *et al*. 2007; Pattyn *et al*. 2021). *ACS* genes are regulated differently based on environmental and developmental factors (Argueso et al., 2007; Dubois et al., 2018). Arabidopsis has eight identified *ACS* genes (Argueso *et al*., 2007). The final step is catalyzed by ACC oxidase (ACO), which converts ACC to ethylene, cyanide, and CO_2_ (Argueso *et al*., 2007; Houben & Van de Poel, 2019). Arabidopsis has around five *ACO* genes (Houben & Van de Poel, 2019), while cucumber has four *ACO* genes involved in sex determination (Chen *et al*., 2016a).

The genes involved in ethylene signaling are highly conserved among plants. Ethylene receptors, including *ethylene response sensor 1* (*ERS1*), *ERS2*, *ethylene resistance 1* (*ETR1*), *ETR2*, and *ethylene insensitive 4* (*EIN4*), found in the endoplasmic reticulum, bind to ethylene to initiate signaling (Dubois *et al*., 2018). Ethylene acts as an inverse agonist, promoting its own production by altering the activity of the biosynthetic pathway and reducing inhibition of downstream signaling (Mayerhofer *et al*., 2012; Binder, 2020). Ethylene inhibits *constitutive triple response 1* (*CTR1*) activity, leading to dephosphorylation of EIN2 and cleavage of its c-terminal domain (C-END). This allows for the repression of *EBF1/2* translation by binding to the 3’ untranslated regions of their mRNA (Chang, 2016; Dubois *et al*., 2018; Binder, 2020). In the absence of ethylene, *EBF1/2* target the primary ethylene-responsive transcription factors *EIN3* and *EIN3-LIKE* (*EIL1*) for degradation (Dubois *et al*., 2018). Repression of *EBF1/2* by ethylene enables the expression of *EIN3* and *EIN3-like* transcription factors, which induce the expression of ethylene response factors (ERFs), further promoting ethylene biosynthesis. Although the canonical signaling pathways are well understood, further investigation is needed regarding ERFs (Olsen *et al*., 2015; Dubois *et al*., 2018).

Compared with dioecious cannabis, the role of ethylene in sex determination in monoecious members of the Cucurbitaceae family has been much more well-defined. In cucurbits ethylene biosynthesis and signaling have also been shown to be manipulated by STS or ethephon applications (Li *et al*., 2021). Typically, ethylene (usually applied as ethephon) has a feminizing effect, increasing the ratio of female flowers relative to males, while the use of STS has the inverse effect, causing a masculinizing effect and results in increased production of male flowers. Numerous studies of sex determination in cucurbits have pointed to the role of ethylene biosynthesis genes such as *ACS4* and *ACS11*, as well as ethylene signaling genes such as the ethylene receptor *ETR1* as contributing to sex determination (Boualem *et al*., 2015; Zhang *et al*., 2017; Li *et al*., 2020). This has led to the labelling of some key loci in Cucurbitaceae as sex-determining loci, with changes of expression of these particular loci determining the sex of the plant, in contrast to *C. sativa* where the chromosome karyotype determines plant sex. The manipulation of cucumber sex has allowed growers to control and optimize the ratio of female to male flowers to improve yield and to control production so as to meet labour constraints (Oda *et al*., 2022). In cannabis, control of plant sex is important as medicinally valuable cannabinoids are primarily produced in female flowers. However, without a pathway model for ethylene biosynthesis and signaling in the species, study of cannabis sexual plasticity remains impossible. The highly conserved nature of ethylene biosynthesis and signaling and well-characterized pathways in model species *Arabidopsis thaliana* present a valuable starting point for the reconstruction of the pathway in an under-researched plant such as *C. sativa* where ethylene biosynthesis and signaling has yet to be explored. In the current study, we use homology analysis and RNA-seq data to present the first *C. sativa* putative Yang cycle, ethylene biosynthesis and ethylene signaling pathways, which will serve as an important foundation for expanding our understanding of the role of ethylene in sexual plasticity of this under-researched dioecious species.

## Materials & Methods

### *In silico* identification of ethylene-related gene orthologs in *C. sativa*

*Arabidopsis thaliana* was chosen as the model from which a list of genes related to ethylene biosynthesis and signaling, as well as the production of ethylene precursors derived from the Yang cycle were compiled following a review of the literature (Pommerrenig *et al*., 2011; van de Poel *et al*., 2012; Olsen *et al*., 2015; Chang, 2016; Pattyn *et al*., 2021). The resulting list of *A. thaliana* ethylene-related genes (AtERGs; Supplementary Table 2) were used for the identification of orthologous *C. sativa* ERGs (CsERGs). Nucleotide and protein sequences for AtERGs were obtained from the National Center for Biotechnology Information database (NCBI; Bethesda, MD). Ortholog identification was conducted on the *C. sativa* cs10 reference genome (RefSeq assembly accession: GCF_900626175.2; Grassa *et al*., 2021). A list of putative CsERG orthologs was compiled using two different methods. The first method employed OrthoFinder (Emms & Kelly, 2015) adapted from Carey *et al*. (2021) and the second method consisted of a BLASTP search of each individual AtERG against the cs10 reference genome (Altschul *et al*., 1990), followed by a manual selection of putative orthologs from the of BLASTP result adapted from Barcaccia *et al*. (2020). This method is described in further detail in the supporting information. The chromosome map (Figure 4a) was generated with chromoMap (v4.1.1; Anand & Rodriguez Lopez, 2022) and bar plots of gene distribution (Figure 4b) was generated with ggplot2 (v3.4.1; Wickham, 2016) using the colour palette Spectral from RColorBrewer (Brewer *et al*., 2002). The protein sequences of the putative CsERGs were then analysed using the InterProScan tool on the InterPro (v97.0; Paysan-Lafosse *et al*., 2023) database to identify conserved protein function or conserved domains and compared with those present in AtERG orthologs (**Supplementary Table *3*** Results of an InterProScan of the CsERGs. Protein family is given when available, otherwise the conserved protein domains are provided, alongside the InterPro entry for each family/domain.Supplementary Table 3).

### RNA-seq data collection and processing

The RNA-seq reads from Adal *et al*. (2021) and Prentout *et al*. (2020), hereafter referred to as dataset 1 and dataset 2, were downloaded from the Sequence Read Archive (SRA) on NCBI under the BioProject accessions PRJNA669389 and PRJNA549804, respectively. RNA was isolated from male flowers (MFs), female flowers (FFs) and STS-induced IMFs for dataset 1, and from MFs and FFs for dataset 2. To perform a relevant comparison between the 2 datasets, only data from similar tissues were considered, i.e., the young flower buds. The SRA sample accessions used in this study are indicated in Supplementary Table 1. FastQC (Andrews, 2010) was used for validating the quality of the data. Then, reads were aligned to the *C. sativa* cs10 reference genome (version GCF_900626175.2; Grassa *et al*., 2021) using STAR (v2.7.2b; Dobin *et al*., 2013) as reported in Cai *et al*. (2020). The STAR index was built with the –sjdbOverhang=100 and the genomeSAindexbases=13 arguments. Splice junctions from the *C. sativa* cs10 RefSeq assembly GTF version GCF_900626175.2 were provided via the --sjdbGTFfile argument. STAR alignments were generated using the default parameters including the --outSAMtype BAM SortedByCoordinate arguments. Quality control of the mapped reads was performed using Qualimap (v2.2.1; Okonechnikov *et al*., 2015). One IMF sample from dataset 1 was excluded based on abnormal distribution of GC-content among mapped reads which included large shoulders at 30% and 60 % GC content, indicating contamination of the sample (see Supplementary Table 1). Finally, quantification of reads overlapping genetic features was performed with HTSeq (v2.0.1; Putri *et al*., 2022) based on the cs10 GFF file obtained from NCBI (https://ftp.ncbi.nlm.nih.gov/genomes/all/GCF/900/626/175/GCF_900626175.2_cs10/GCF_900626175.2_cs10_genomic.gff.gz) in order to generate a gene quantification table required for subsequent differential analysis with DEBrowser (v1.22.5; Kucukural *et al*., 2019).

### Differential expression analysis of CsERGs

Differentially expressed CsERGs were identified using DESeq2 (Love *et al*., 2014) with an FDR-adjusted *p*-value < 0.05 on the DEBrowser graphical user interface in RStudio (RStudio Inc., MA, USA) using v4.1.1 of R. The analysis was adapted from DEBrowser’s DESeq2 method outlined on the DEBrowser Github (https://github.com/UMMS-Biocore/debrowser). In short, gene quantification tables generated in HTSeq and a manually generated metadata file for the samples were uploaded to DEBrowser. Genes with count per million values less than 1 (CPM < 1) in 7 out of 15 replicates in dataset 1 and 6 out of 12 replicates in dataset 2 were removed. Replicates were then grouped and pairwise comparisons were performed based on the floral type. In DEBrowser the following DESeq2 parameters were used following the method outline by Kucukural *et al*. (2019) and the DEBrowser Github: *Fit Type-parametric; betaPrior-FALSE; Test Type-Wald; Shrinkage-None*. Differential expression levels were then normalized using a Median Ratio Normalization (MRN) and pairwise comparisons between treatments were considered significantly different in subsequent comparisons if they had an adjusted *p*-value <0.05 according to the Wald test and a |log2foldchange| > 1. Heatmaps comparing differential gene expression were then generated using DEBrowser heatmap function. Tables from pairwise comparisons were generated from DEBrowser and then used for the generation of gene expression bar plots using ggplot2 (Wickham, 2016) in RStudio.

## Results

### ERGs are conserved in *C. sativa*

Based on 47 AtERGs previously identified (Supplementary Table 2), ortholog analyses identified 43 putative CsERGs (Table 1). At least one ortholog of each key gene in the canonical ethylene biosynthesis and signaling pathways, as well as the Yang cycle, was identified *in silico* except for auxin-regulated genes involved in organ size (*AtARGOS*; Supplementary Table 2). Analysis of the amino acid sequences of the CsERGs also confirmed the presence of most conserved domains found in the AtERGs (see Supplementary Table 3). To the best of our knowledge, this is the first time that the genes coding for the various enzymes involved in the putative Yang cycle (Figure 2a), ethylene biosynthesis (Figure 2b) and signaling pathway (Figure 3) have been proposed in *C. sativa.* These genes are available in a collection on the NCBI site at: https://www.ncbi.nlm.nih.gov/sites/myncbi/adrian.monthony.1/collections/63471644/public/.

**Figure 2.**
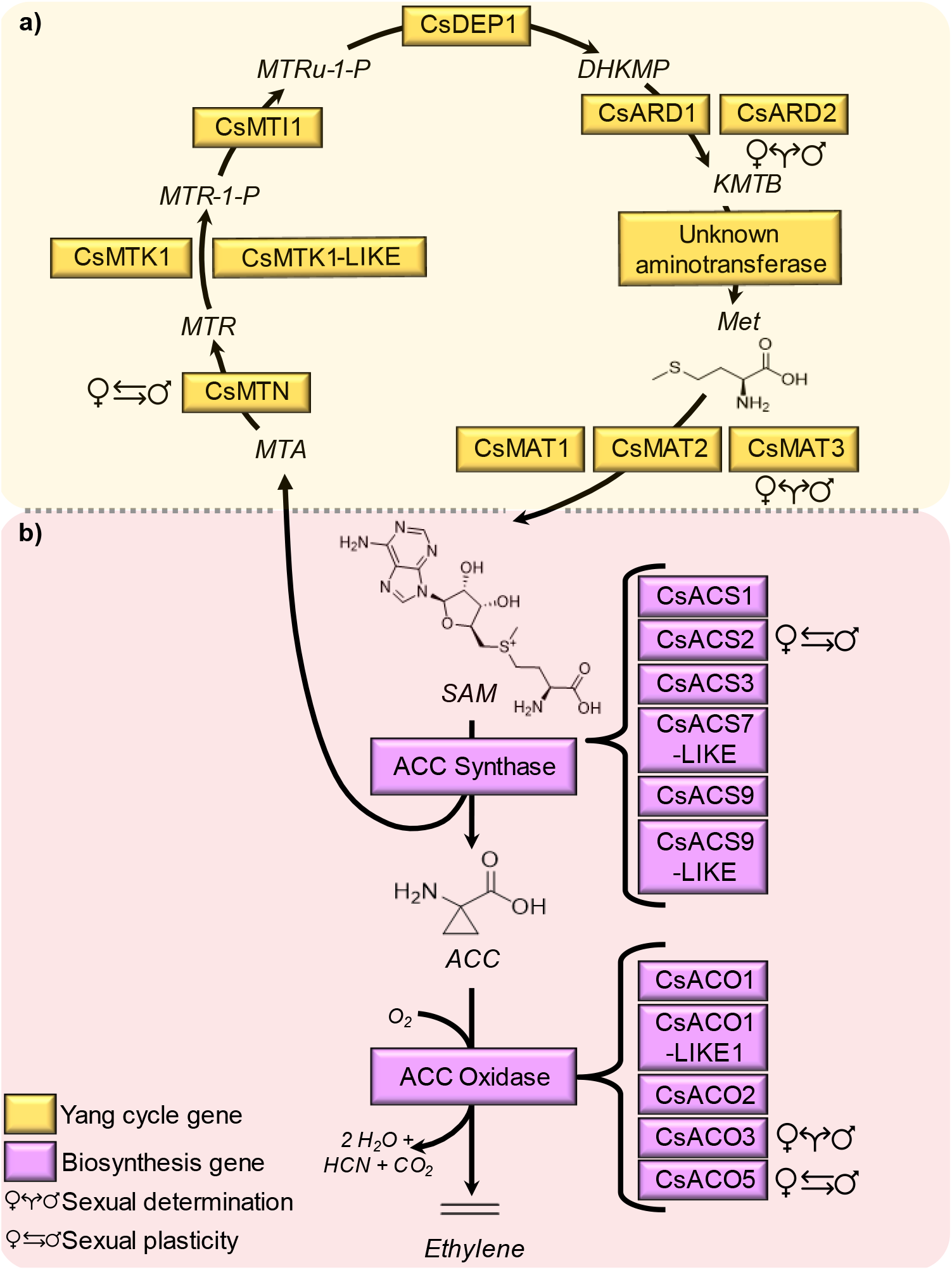
*C. sativa* model of the Yang cycle and ethylene biosynthesis determined by homology analysis from *A. thaliana.* Products of enzymatic reactions are italicized. Genes encoding enzymes are in boxes. (a) Yang cycle-product abbreviations are as follows: *S-* adenosylmethionine (SAM), 5-methylthioadenosine (MTA), 5-methylthioribose (MTR), 5-methyltioriobose-1-P (MTR-1-P), 5-methylthioribulose-1-P (MTRu-1-P), 1,2-dihydroxy-3-keto-5-methylthiopentene (DHKMP), 2-keto-4-methylthiobutyrate (KMTB), methionine (Met). (b) Ethylene biosynthesis-product abbreviations are as follows: 1-aminocyclopropane-1-carboxilic acid (ACC). Differentially expressed genes are annotated with Mars and Venus symbols, identifying their proposed roles as either involved in sexual determinism (sex determination; denoted with diverging arrows) or sexual plasticity (denoted with reciprocal arrows) based on RNA-seq analysis.

**Figure 3.**
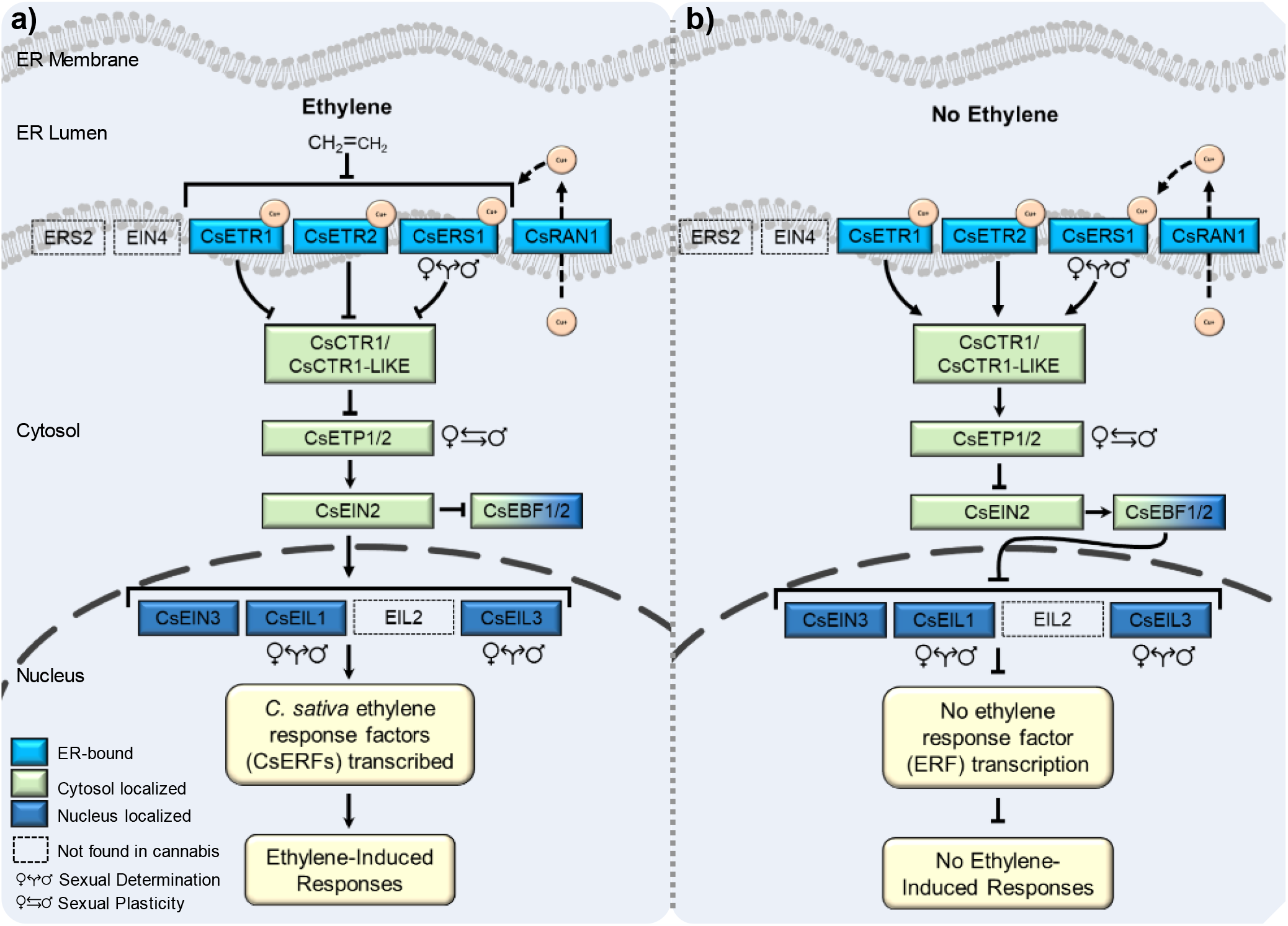
A simplified model of ethylene signaling in *C. sativa*, constructed through computational analysis using the *A. thaliana* pathway as a template. Pathway is depicted in the presence (a) and absence-(b) of ethylene. Dashed lines represent genes not found in *C. sativa*. Pointed arrows indicate promotion, while blunt-end arrows represent repression. The figure design is based on existing ethylene signaling models (Binder *et al*., 2010; Dubois *et al*., 2018; Binder, 2020). Differentially expressed genes are marked with Mars and Venus symbols, indicating their proposed roles in sexual determinism (sex determination, diverging arrows) or sexual plasticity (reciprocal arrows) based on RNA-seq analysis.

**Figure 4.**
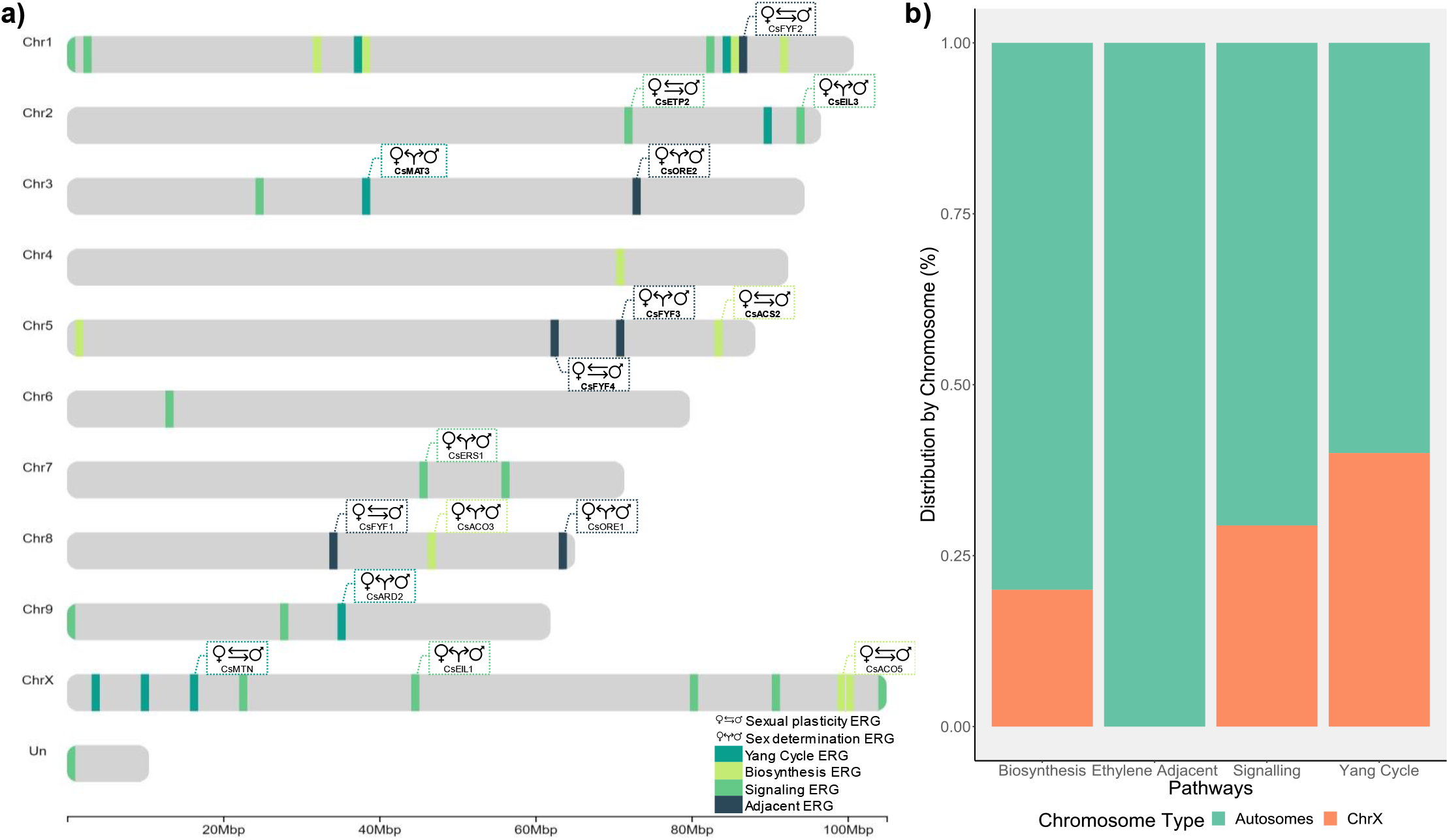
Position of CsERGs in the cs10 reference genome identified from A. thaliana orthologs. (a) Distribution of CsERGs across the genome. Distinct colors were used to describe their functional annotation. (b) Proportion of CsERGs distributed on autosomes or ChrX. An interactive version of (a) can be found in the on the OSF page for this project.

**Table 1.** *Cannabis sativa* ethylene-related genes (CsERGs) identified through ortholog analysis.

| Cs10 ERG Locus* | Name | Chromosome | *A. thaliana*<br>orthologs | Pathway | Identification<br>method |
| --- | --- | --- | --- | --- | --- |
| LOC115704991 | *CsACO1* | 1 | *AT2G19590* | Biosynthesis | BLASTP,<br>OrthoFinder |
| LOC115702109 | *CsACO1-*<br>*LIKE1* | Un<sup>§</sup> | *AT2G19590* | Biosynthesis | BLASTP,<br>OrthoFinder |
| LOC115707397 | *CsACO2*<sup>*†</sup> | 1 | *AT1G62380*,<br>*AT1G12010*,<br>*AT1G05010* | Biosynthesis | BLASTP,<br>OrthoFinder |
| LOC115699412 | *CsACO3*<sup>*†</sup> | 8 | *AT1G62380*,<br>*AT1G12010*,<br>*AT1G05010* | Biosynthesis | BLASTP,<br>OrthoFinder |
| LOC115699408 | *CsACO5*<sup>*†</sup> | X | *AT1G77330* | Biosynthesis | BLASTP,<br>OrthoFinder |
| LOC115704849 | *CsACS1* | 1 | *AT3G49700*,<br>*AT4G08040*,<br>*AT4G11280*,<br>*AT1G01480* | Biosynthesis | BLASTP |
| LOC115717044 | *CsACS2*<sup>*†</sup> | 5 | *AT4G08040*,<br>*AT1G01480*,<br>*AT4G11280*,<br>*AT3G49700* | Biosynthesis | BLASTP,<br>OrthoFinder |
| LOC115712396 | *CsACS3* | 4 | *AT5G65800*,<br>*AT4G37770*,<br>*AT4G08040*,<br>*AT3G49700* | Biosynthesis | BLASTP,<br>OrthoFinder |
| LOC115704963 | *CsACS7-*<br>*LIKE* | 1 | *AT4G26200*,<br>*AT1G01480*,<br>*AT4G11280*,<br>*AT4G08040*,<br>*AT3G49700* | Biosynthesis | BLASTP,<br>OrthoFinder |
| LOC115696400 | *CsACS9*<sup>†</sup> | X | *AT4G08040*,<br>*AT5G65800*,<br>*AT4G37770*,<br>*AT3G49700* | Biosynthesis | BLASTP |
| LOC115716540 | *CsACS9-*<br>*LIKE* | 5 | *AT5G65800*,<br>*AT4G37770*,<br>*AT4G08040*,<br>*AT3G49700*,<br>*AT2G22810* | Biosynthesis | BLASTP |
| LOC115697744 | *CsCTR1*<sup>*†</sup> | X | *AT5G03730* | Signaling | BLASTP |
| LOC115706236 | *CsCTR1-*<br>*LIKE*<sup>*†</sup> | 1 | *AT5G03730* | Signaling | BLASTP,<br>OrthoFinder |
| LOC115720532 | *CsEBF1*<sup>*†</sup> | X | *AT5G25350*,<br>*AT2G25490* | Signaling | BLASTP,<br>OrthoFinder |
| LOC115724585 | *CsEBF2*<sup>*†</sup> | 6 | *AT5G25350*,<br>*AT2G25490* | Signaling | BLASTP |
| LOC115711236 | *CsEIL1*<sup>*†</sup> | X | *AT2G27050* | Signaling | OrthoFinder |
| LOC115719373 | *CsEIL3*<sup>*†</sup> | 2 | *AT1G73730* | Signaling | OrthoFinder,<br>BLASTP |
| LOC115709621 | *CsEIN2*<sup>*†</sup> | 3 | *AT5G03280* | Signaling | BLASTP,<br>OrthoFinder |
| LOC115706079 | *CsEIN3*<sup>*†</sup> | 1 | *AT2G27050*,<br>*AT3G20770* | Signaling | BLASTP,<br>OrthoFinder |
| LOC115723050 | *CsERF1* | 9 | *AT3G23240* | Signaling | BLASTP,<br>OrthoFinder |
| LOC115697291 | *CsERS1*<sup>*†</sup> | 7 | *AT2G40940* | Signaling | BLASTP,<br>OrthoFinder |
| LOC115705474 | *CsETP1*<sup>*†</sup> | 1 | *AT3G18980* | Signaling | BLASTP |
| LOC115720323 | *CsETP2* | 2 | *AT3G18910* | Signaling | BLASTP |
| LOC115702329 | *CsETR1*<sup>*†</sup> | X | *AT1G66340*,<br>*AT2G40940* | Signaling | BLASTP,<br>OrthoFinder |
| LOC115721785 | *CsETR2*<sup>*†</sup> | 9 | *AT3G04580*,<br>*AT1G04310*,<br>*AT3G23150* | Signaling | BLASTP,<br>OrthoFinder |
| LOC115698184 | *CsRAN1*<sup>*†</sup> | 7 | *AT5G44790* | Signaling | BLASTP,<br>OrthoFinder |
| LOC115707134 | *CsRTE1*<sup>*†</sup> | X | *AT2G26070* | Signaling | BLASTP,<br>OrthoFinder |
| LOC115721731 | *CsARD1*<sup>*†</sup> | 9 | *AT5G43850*,<br>*AT4G14716*,<br>*AT4G14710*,<br>*AT2G26400* | Yang Cycle | BLASTP,<br>OrthoFinder |
| LOC115722194 | *CsARD2*<sup>*†</sup> | 9 | *AT4G14716*,<br>*AT2G26400*, | Yang Cycle | BLASTP,<br>OrthoFinder |
|  |  |  | *AT5G43850*,<br>*AT4G14710* |  |  |
| LOC115706479 | *CsMTI1*<sup>*†</sup> | 1 | *AT2G05830* | Yang Cycle | BLASTP,<br>OrthoFinder |
| LOC115703014 | *CsMTK1*<sup>*†</sup> | X | *AT1G49820* | Yang Cycle | BLASTP,<br>OrthoFinder |
| LOC115702983 | *CsMTK1-*<br>*LIKE*<sup>*†</sup> | X | *AT1G49820* | Yang Cycle | BLASTP,<br>OrthoFinder |
| LOC115704327 | *CsMTN*<sup>*†</sup> | X | *AT4G34840*,<br>*AT4G38800* | Yang Cycle | BLASTP,<br>OrthoFinder |
| LOC115706293 | *CsDEP1*<sup>*†</sup> | 1 | *AT5G53850* | Yang Cycle | BLASTP,<br>OrthoFinder |
| LOC115711364 | *CsMAT1*<sup>*†</sup> | X | *AT1G02500*,<br>*AT4G01850*,<br>*AT3G17390*,<br>*AT2G36880* | Yang Cycle | BLASTP,<br>OrthoFinder |
| LOC115719259 | *CsMAT2*<sup>*†</sup> | 2 | *AT1G02500*,<br>*AT4G01850*,<br>*AT3G17390*,<br>*AT2G36880* | Yang Cycle | BLASTP |
| LOC115711343 | *CsMAT3*<sup>*†</sup> | 3 | *AT2G36880*,<br>*AT3G17390*,<br>*AT4G01850*,<br>*AT1G02500* | Yang Cycle | BLASTP,<br>OrthoFinder |
| LOC115700576 | *CsFYF1*<sup>*†</sup> | 8 | *AT5G62165* | Ethylene<br>Adjacent | BLASTP |
| LOC115706939 | *CsFYF2*<sup>*†</sup> | 1 | *AT5G62165* | Ethylene<br>Adjacent | BLASTP |
| LOC115716986 | *CsFYF3*<sup>*†</sup> | 5 | *AT5G62165* | Ethylene<br>Adjacent | BLASTP |
| LOC115717309 | *CsFYF4*<sup>*†</sup> | 5 | *AT5G62165* | Ethylene<br>Adjacent | BLASTP |
| LOC115701489 | *CsORE1*<sup>*†</sup> | 8 | *AT5G39610* | Ethylene<br>Adjacent | BLASTP,<br>OrthoFinder |
| LOC115709772 | *CsORE2*<sup>*†</sup> | 3 | *AT5G39610* | Ethylene<br>Adjacent | BLASTP |

\* Hyperlinks to the NCBI cs10 reference genome information page for the specific locus.
\* and † ERGs for which transcripts were detected from dataset 1 and/or in dataset 2, respectively.
§ A gene located on an unclassified scaffold.

### Differential expression analysis reveals three patterns of expression

Among the 43 putative CsERGs analyzed, a total of 35 were found to be expressed in both dataset 1 and dataset 2, as indicated in Table 1. Upon further investigation, we found that among these 35 CsERGs, 16 exhibited differential expression between flower types in dataset 1, while 11 showed differential expression in dataset 2, as shown in Supplemental Figure 2. Seven CsERGs that consistently displayed differential expression between MFs and FFs were identified across both datasets, with comparable levels of magnitude, as illustrated in Supplemental Figure 3. These 7 CsERGs are considered high confidence genes involved in cannabis sex determination or sexual plasticity. In conducting pairwise comparisons of CsERG expression between different flower types, we observed three distinct patterns of differential expression. These patterns are described below and represent a newly proposed framework for understanding the role of CsERGs in sex determination and sexual plasticity. These patterns have been classified as karyotype concordant (KC-sex determination; Figure 5), floral organ concordant (FOC-sexual plasticity; Figure 6), and unique ERG (uERG-sexual plasticity; Figure 7).

**Figure 5.**
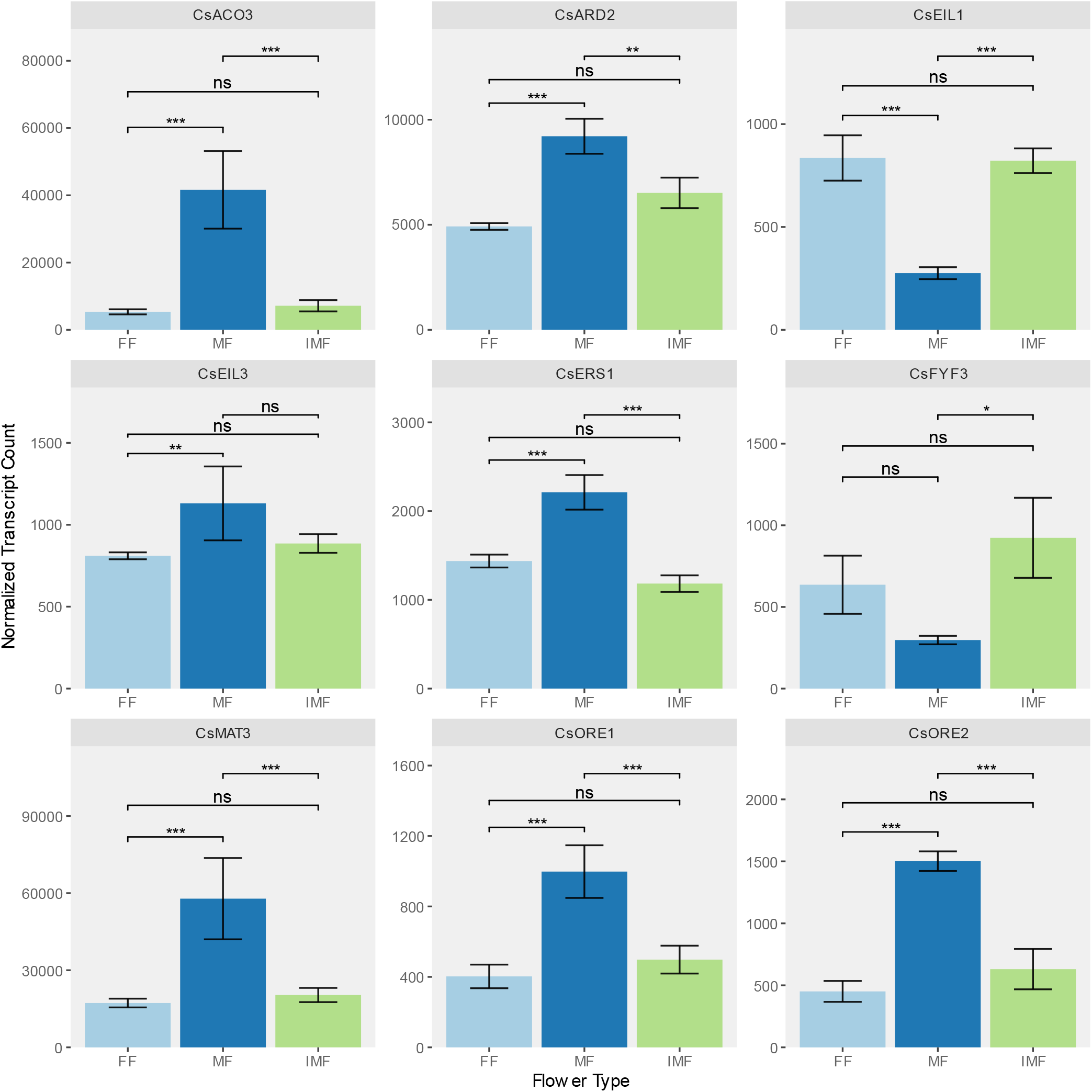
Differential expression analysis of *C. sativa* ERGs revealed nine genes with a karyotype concordant (KC) expression pattern. Among these genes, female flowers (FF; XX karyotype) and induced male flowers (IMF; XX karyotype) showed not significant (ns) differential expression. Significant pairwise comparisons are denoted by asterisks: ns = not significant; * for p ≤ 0.05; ** for p ≤ 0.01; *** for ≤ 0.001 as determined by the Wald test.

**Figure 6.**
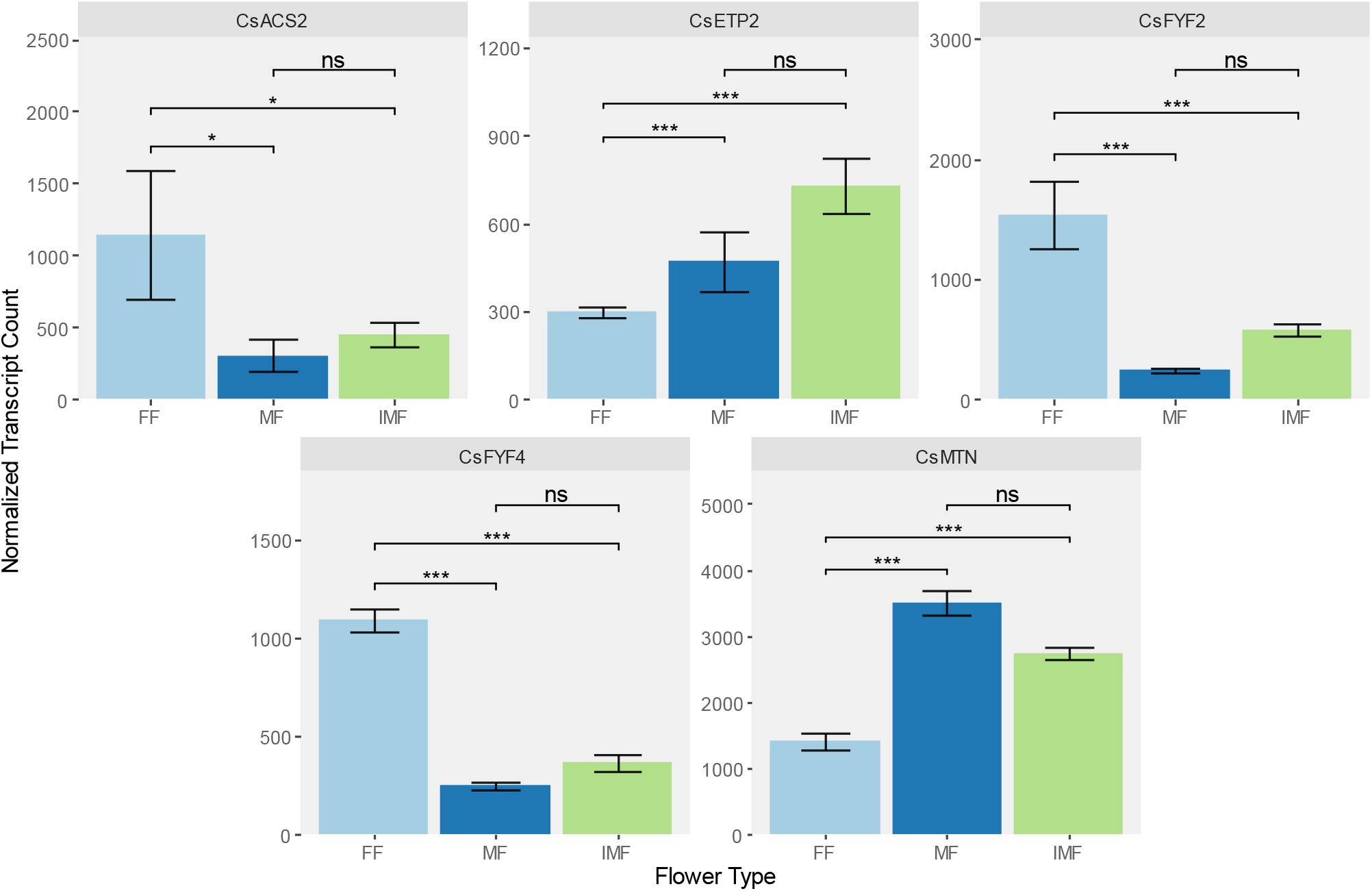
Differential expression analysis of *C. sativa* ERGs revealed five cases of genes showing floral-organ concordant (FOC) expression. FOC refer to genes whose expression pattern is shared between organisms with the same floral organ, but not necessarily the same sex-chromosome karyotype designated by ns in transcription levels for male flowers on XY plants and induced male flowers on XX plants. S0 Significant pairwise comparisons are denoted by asterisks: ns = not significant; * for p ≤ 0.05; ** for p ≤ 0.01; *** for ≤ 0.001 as determined by the Wald test.

**Figure 7.**
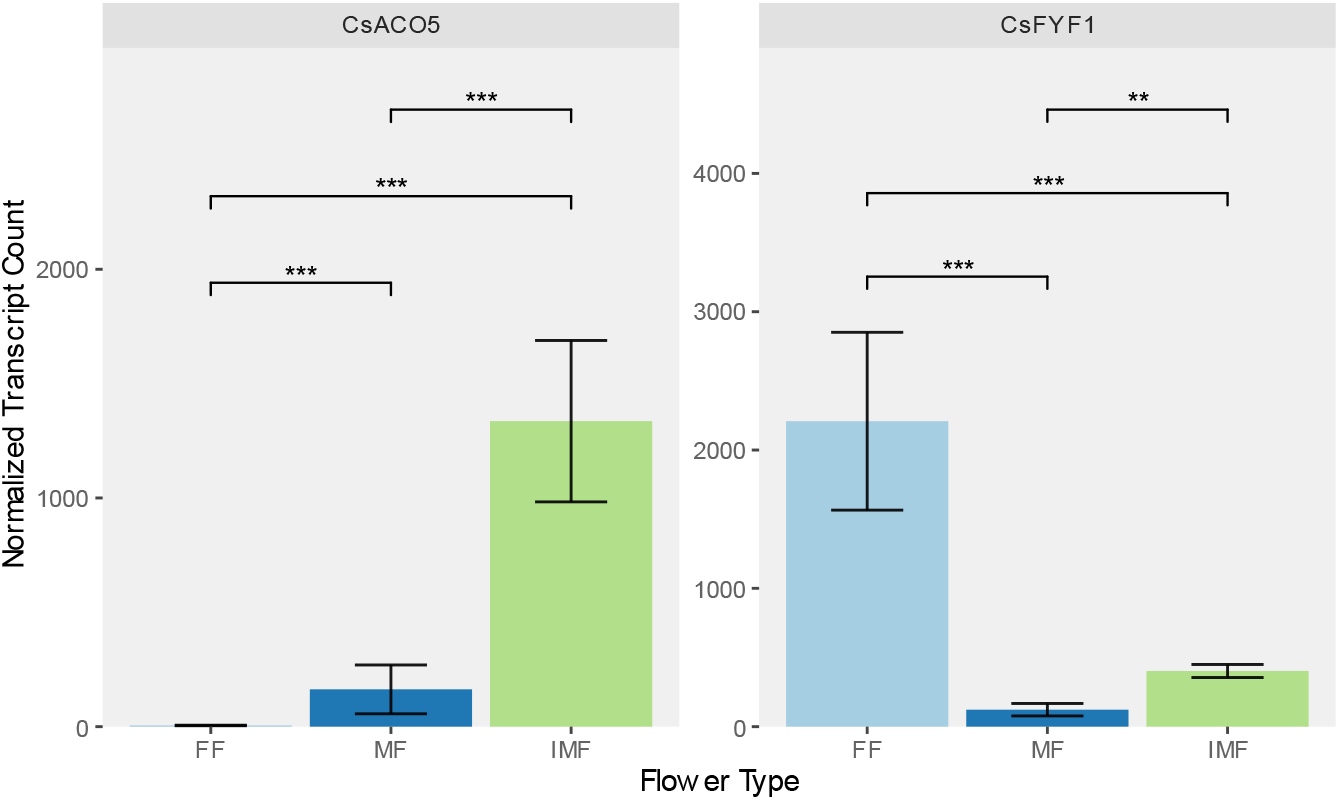
Differential expression analysis of revealed unique ERG (uERG) expression, which are cases where gene expression in the sex-changed plant does not match either that of the plant with which is shares the same sex chromosome karyotype or the plant with which is shares the same floral organs. Significant pairwise comparisons are denoted by asterisks: ns = not significant; * for p ≤ 0.05; ** for p ≤ 0.01; *** for ≤ 0.001 as determined by the Wald test.

KC expression refers to CsERGs that displayed no significant differences in expression between plants with the same sex chromosome karyotype, such as FFs and IMFs. Among the 16 CsERGs exhibiting differential expression, 9 were identified to have KC expression, primarily in dataset 1, as depicted in Figure 5. Conversely, FOC expression describes genes with similar expression between MFs and IMFs, which possess the same floral organs, but that had significantly different expression when compared with FF. Out of the 16 differentially expressed CsERGs, five displayed FOC expression, as indicated in Figure 6. Notably, three genes (*CsACS2, CsFYF2*, and *CsFYF4*) exhibited higher expression in FFs compared to phenotypic male flowers, whereas two genes (*CsETP2* and *CsMTN*) demonstrated higher expression in MFs and IMFs compared to FFs, as illustrated in Figure 6. Lastly, uERG expression referred to CsERGs that exhibited significant differential expression among each floral type, including FFs, MFs, and IMFs. Two CsERGs, namely *CsACO5* and *CsFYF1*, were identified to possess uERG expression, as depicted in Figure 7. By proposing this framework, we aim to deepen our understanding of the involvement of CsERGs in sex determination and sexual plasticity in cannabis.

### ERGs are distributed genome-wide in *C. sativa*

The 43 CsERGs identified in the homology analysis were distributed across the genome (Figure 4 and Supplementary Table 4), independently of autosomes or sex chromosomes. Approximately 74% (32) of the CsERGs were located on autosomes (Figure 4b). Yang cycle genes constituted the majority (40%; 3) of CsERGs on the X chromosome. In contrast, none of the genes that were classified as ethylene adjacent (ERGs from the literature not belonging to the canonical ethylene biosynthesis or signaling pathways) were found on the X chromosome (Figure 4). Among the 43 CsERGs identified, all but one (*CsACO1-LIKE1*) mapped to chromosomes in the reference genome, with *CsACO1-LIKE1* located on an unassigned scaffold (Figure 4a). CsERGs exhibiting significant differential expression associated with sex determination (KC) and sexual plasticity (FOC and uERG) were distributed throughout the genome (Figure 4a), encompassing genes from the Yang Cycle and ethylene biosynthesis pathway (Figure 2) as well as the ethylene signaling pathway (Figure 3). Approximately 37% of the ERGs (16) displayed a pattern of differential expression, among which only three were mapped to the sex chromosomes (Figure 4a). Specifically, *CsEIL1* exhibited KC expression patterns, while *CsACO5* and *CsMTN* showed FOC/uERG expression. The remaining 13 differentially expressed genes were located on various autosomes (Figure 4a), except for chromosomes 4 and 6, which lacked any differentially expressed ERGs.

## Discussion

To date, sexual plasticity has only been discussed in the context of the animal kingdom (Liu *et al*., 2017). While studies have focused on sex determination in angiosperms without sex chromosomes, such as the Cucurbitaceae family (Yamasaki *et al*., 2001; Boualem *et al*., 2015; Oda *et al*., 2022), sexual plasticity in plants remains relatively unexplored. However, as a dioecious plant with sex chromosomes, cannabis represents an uncommon example of a plant species that exhibits sexual plasticity. Previous research has demonstrated the induction of male flowers in female plants through the inhibition of ethylene using STS (Lubell & Brand, 2018), and the induction of female flowers in male plants through ethylene (ethephon) treatment (Moon *et al*., 2020a). Considering ethylene’s known involvement in sex determination across plant species, it serves as a promising starting point for investigating sexual plasticity in dioecious plants like *C. sativa* (Hume & Lovell, 1983; Manzano *et al*., 2014; Cebrián *et al*., 2022). However, unraveling the specific mechanisms by which ethylene influences sexual plasticity presents challenges due to its multifaceted roles in plants.

To understand the influence of CsERGs on sex determination and sexual plasticity, we propose a framework consisting of three putative CsERG expression patterns: karyotype concordant (KC; Figure 5), floral organ concordant (FOC; Figure 6), and unique ERG (uERG; Figure 7). KC expression is similar between individuals with the same sex chromosome karyotype and CsERGs exhibiting KC expression are unaffected by STS treatments that induce sexual plasticity in *C. sativa*. This type of expression suggests that genes belonging to the KC case may have sex-specific patterns of expression, playing a role in sex determination. Conversely, FOC expression (Figure 6) is the same between individuals with shared floral organ sex, but carrying different sex chromosome karyotypes. This pattern suggests that CsERGs showing FOC expression are involved in sexual plasticity, as their expression appears uncoupled from the sex chromosome karyotypes. uERGs (Figure 7) shows a pattern of distinct levels of expression of CsERGs in all three treatments. Two genes, *CsFYF1* and *CsACO5*, showed uERG expression, with the gene expression in the sex-manipulated plant (IMF) not matching either of the plants with the same genotypic sex (FF) or the plant with the phenotypic sex (MF). Below, we discuss how each of these patterns is found throughout the biochemical journey of ethylene, from its precursor producing in the Yang cycle (Figure 2a) all the way through ethylene biosynthesis (Figure 2b) and signaling (Figure 3). We also highlight that the distribution of these differentially expressed CsERGs is genome-wide and not exclusive to the sex chromosomes (Figure 4). Lastly, we end by discussing limitations and future opportunities revealed by this study.

### High expression of key Yang Cycles genes; a potential indicator of masculinity in *C. sativa*

In previous studies, it has been demonstrated that changes in the expression of Yang cycle genes regulate ethylene production in plants. Here, we explore the potential role of the Yang cycle in cannabis sex determination and sexual plasticity. In the present study, we found that two CsERGs from the Yang cycle showed KC expression (Figure 2a and Figure 5), methionine adenosyltransferase 3 (*CsMAT3*) and acireductone dioxygenase 2 (*CsARD2*). Most higher eukaryotes possess small *ARD* gene families (Sauter *et al*., 2005). *A. thaliana* has four *ARD* genes isoforms (Pommerrenig *et al*., 2011) from which we identified two *CsARD* genes homologs. In the dataset 1, *CsARD2* was more highly expressed in MFs than FFs and IMFs (Figure 5). In contrast *CsARD1* transcripts were found in all treatments, in both datasets, but expression did not vary significantly between MFs, FFs or IMFs. These findings are consistent with previous studies in rice that showed that the different ARD isoforms are regulated differently *in planta*, with *OsARD1* undergoing a transient increase in expression in response to submergence, while *OsARD2* is constitutively expressed, and unaffected by changes in ethylene concentration (Sauter *et al*., 2005). Similarly, Liang *et al*. (2019) showed that transgenic rice overexpressing *OsARD1* produced higher levels of ethylene versus wild-type rice.

Methionine adenosyltransferase (MAT, or SAM synthetase; SAMS) is responsible for the biosynthesis of SAM, an important methyl donor and precursor for ethylene, lignin and polyamine synthesis (Arraes *et al*., 2015). From the four *AtMAT* used to construct the putative Yang cycle model in cannabis, we identified three putative *CsMAT* genes: *CsMAT1, CsMAT2, CsMAT3,* of which *CsMAT3* was found to have a KC expression. *CsMAT3* was most expressed in MFs, with significantly lower transcription levels in FFs and IMFs. The association *MAT* genes with male floral characteristics have been reported in *Arabidopsis* by Chen *et al*. (2016b) who found that expression of *AtMAT3,* a *CsMAT3* ortholog, is responsible for normal pollen germination and pollen tube formation. Although critical male fertility and pollen germination in *Arabidopsis*, the role of *MAT* genes in controlling sex determination has yet to be reported. The present findings that *CsMAT3* gene expression is higher in MFs than in FFs, suggest that it may play a role in sex determination. One way that *CsMAT3* may affect sex determination is by altering levels of DNA methylation. SAM, the product of *MAT* is a methyl donor for DNA and histone methylation, and it has been reported that *mat4* knockouts in *Arabidopsis* have lower genome-wide methylation levels as well as decreased histone methylation (Meng *et al*., 2018). Similar findings have been echoed by Chen *et al*. (2016b) who reported that *MAT3* was required for proper histone and tRNA methylation. It is possible that higher expression of *CsMAT3* in male plants could lead to increased methylation of key genes involved in plant masculinity. To date, there has been a scarcity of research on the methylation patterns of cannabis (Aina *et al*., 2004; Mayer *et al*., 2015), with no studies investigating its role in sex determination and sexual plasticity. Investigating the methylation status of floral tissues in cannabis may yield valuable insights into the mechanisms underlying sex determination and sexual plasticity in this species.

Here, we report the identification of one CsERG from the Yang cycle, *CsMTN,* with a FOC expression. MTN recycles MTA after the first step of the ethylene biosynthesis cycle where SAM is converted to ACC by ACS. The ortholog analysis for both *AtMTN1* and *AtMTN2* revealed a single ortholog in cannabis (Table 1), which is similar to both rice (*Oryza sativa* L.) and tomato (*Solanum lycopersicum* L. Karst. ex Farw) in which MTN is only encoded by one gene (Rzewuski *et al*., 2007; van de Poel *et al*., 2012). In Arabidopsis, *MTN* shows preferential expression in the phloem (Pommerrenig *et al*., 2011), suggesting that tissue sampling may also play a role in the expression patterns of *MTN*. In dataset 1 *CsMTN* transcripts were most abundant in MFs (Supplemental Figure 3), and in dataset 2 (Figure 6) *CsMTN* transcripts were significantly more expressed in MFs and IMFs than in FFs, suggesting that high abundance of *CsMTN* may be associated with masculinity and sexual plasticity in cannabis.

### Multiple biosynthesis genes linked to sexual plasticity

Ethylene biosynthesis is regulated by a set of conserved enzymes that are found in all plant species. The key enzyme in this two-step process is ACS, which converts SAM, derived from the Yang cycle, into ACC. ACC is then converted into ethylene by the enzyme ACO. These enzymes are conserved across all plant species, and their regulation is tightly controlled by various factors such as hormonal and environmental cues (Bleecker & Kende, 2000). The ortholog analysis found that *ACS* and *ACO* belong to multi-gene families in cannabis, similar to Arabidopsis. In *C. sativa*, 6 *ACS* and 5 *ACO* gene homologs were identified from the 8 *ACS* and 5 *ACO* genes in *A. thaliana* respectively (Figure 2 and Table 1). The presence of *ACS* and *ACO* are critical to ethylene biosynthesis, but the number of genes in each family can vary between species. For example, in *Pyrus ussuriensis* L. (pear) 13 *ACS* and *11 ACO* genes have been identified (Yuan *et al*., 2020), in *Zea mays* L. 5 *ACS* and 15 *ACO* genes have been identified, in rice 6 *ACS* and 9 *ACO* genes, and in tomato there are 9 *ACS* and 7 *ACO* genes (Houben & Van de Poel, 2019; Wu *et al*., 2022). The importance of having multiple functionally redundant ethylene biosynthesis genes has been demonstrated in multiple plants, with studies showing *ACS* and *ACO* genes are expressed at different levels and in different tissues as a function of the plant organ’s needs and developmental stage (van de Poel *et al*., 2012; Van de Poel *et al*., 2014). In ripening tomatoes Van de Poel *et al*. (2012) reported a higher level of *ACS6, ACO2* and *ACO4* expression during baseline (system 1) ethylene production during immature fruit development prior to ripening, but that auto-catalytic increases in ethylene during ripening (system 2) were correlated with increases in *ACS2*, *ACS4*, *ACO3* and *ACO5* expression, while abundance of *ACO2* and *ACO4* transcripts decreased at that same stage. The induction/repression of specific *ACS* genes has also been shown to play a role in sex determination of some *Cucurbitaceae* species (Manzano *et al*., 2014; Boualem *et al*., 2015; Chen *et al*., 2016a; Zhang *et al*., 2017). One study found that expression of *ACS11* was only observed in the vascular bundles of female flowers of monoecious and in hermaphroditic flowers of andromonoecious plants (plants producing male and hermaphrodite flowers), pointing to the role of *ACS11* in carpel development (Boualem *et al*., 2015). Changes in sex determination have also been shown in cucumber and melon by way of disruption of a specific *ACO* genes (*ACO2* in cucumber; *ACO3* in melon) resulting in decreased ethylene production in the carpel region of flowers and causing a shift from a monoecy to androecy resulting in unisexual flower production in cucumber and melons (Chen *et al*., 2016a). The existence of multiple, functionally redundant *ACS* and *ACO* genes, combined with their spatial and temporal expression, allows for the fine tuning of ethylene biosynthesis. This gives plants a high level of control over their development, including sex determination. The present study demonstrates evidence of numerous *ACS* and *ACO* genes distributed throughout the genome of *C. sativa* (Table 1 and Figure 4). Notably, the expression patters of *CsACS2* (Figure 6)*, CsACO3* (Figure 5) and *CsACO5* (Figure 7) represent the three distinct cases of gene expression proposed in this study. These findings collectively suggest a diverse range of effects exerted by the ethylene biosynthesis pathway on cannabis sex determination and sexual plasticity.

*CsACS2*, which is part of the *ACS* gene family responsible for the first step of ethylene biosynthesis, shows a FOC expression linking it to sexual plasticity. *ACS* has been demonstrated to catalyze the rate-limiting reaction in most cases of ethylene production (van de Poel *et al*., 2012; Houben & Van de Poel, 2019; Pattyn *et al*., 2021). This gene is downregulated in MFs and IMFs in comparison to FFs. Comparisons in *CsACS2* expression show similar low expression levels between MFs and IMFs despite their different genotypic sex. Since *ACS* is thought to be rate-limiting, and given the association of ethylene with ‘feminization’ of floral phenotypes we hypothesize that the downregulation of *CsACS2* in IMF relative to FF, points to reduction in ethylene biosynthesis *in planta* which may play a role in the induction of male flowers in genetically female plants. *CsACO3* shows KC expression, with activity in MFs much higher for both than in IMF and FFs. Further study which incorporates multiple tissue sources could prove helpful in understanding *CsACO3*’s KC expression and how this plays a putative role in sex determination, as it remains to be seen if this expression pattern is unique to floral tissues or if this KC expression is also present in vegetative tissues of flowering plants prior to and during early floral development. Lastly, *CsACO5* has a unique expression pattern in all treatments (uERG; Figure 7), with a significantly larger expression in IMFs compared with MFs or FFs. Here, dosage compensation could play a role with downstream inhibition of ethylene signaling by STS leading to increased transcription of *ACO* as IMF plants which are genetically female, fail to perceive ethylene levels normally required for female floral development.

### Ethylene signaling is conserved in *C. sativa*

Ethylene signaling has also been highly conserved across the plant kingdom (Chang, 2016). Ethylene binds to a family of receptors known as ETRs, which are found in the membrane of the endoplasmic reticulum of plant cells (Binder, 2020). These receptors are responsible for transducing the ethylene signal to downstream signaling components. The downstream signaling pathway involves a series of conserved components, including *CTR1*, *EIN2*, and *EIN3*, which are responsible for activating the expression of ethylene-responsive genes (Shakeel *et al*., 2013). In the present study the putative ethylene signaling pathway in *C. sativa* was identified based on orthology with Arabidopsis genes (Figure 3). Transcriptomes obtained from MFs, FFs and IMFs revealed CsERGs adjacent to, or directly involved in the signaling pathway which have FOC expression (*CsETP2, CsFYF2* and *CsFYF4*) and uERG (*CsFYF1*). In *Arabidopsis EIN2 TARGETING PROTEIN1* and *EIN2 TARGETING PROTEIN2* (*ETP1/2*) act to suppress ethylene responses in the signaling pathway by carrying out ubiquitinating of *EIN2*, marking *EIN2* degradation (Qiao *et al*., 2009; Binder, 2020). When ethylene is present, *ETP1/2* is supressed, allowing ethylene signaling to proceed (Qiao *et al*., 2009; Yang *et al*., 2015). In the present study, both *CsETP1* and *CsETP2* transcripts were detected, however only *CsETP2* had an FOC expression associated with sexual plasticity (Figure 3). *CsETP2* expression was lowest in FFs compared to MFs and IMFs (Figure 6). There was no significant difference between *CsETP2* expression in MFs and IMFs and both had higher expression than FF tissues. These patterns of gene expression fit with the hypothesized role of ethylene as a feminizing agent. If increased endogenous ethylene is responsible for femaleness, then we expect lower levels of *CsETP2* in female flower producing plants, as observed.

The differential expression analysis also revealed three *FOREVER YOUNG FLOWER* (*FYF*) genes which belong to the FOC expression (*CsFYF2* and *CsFYF4*) and uERG (*CsFYF1*) expression. *FYF* genes are expressed in young flowers prior to pollination (Chen *et al*., 2011) and acts as a suppressor of floral abscission. They act by repressing *Ethylene Response DNA-binding Factors (EDFs)* found downstream *ERFs* that trigger floral senescence and abscission in maturing flowers (Chen *et al*., 2015, 2022). In both datasets, expression of *CsFYF1* and *CsFYF2* was highest in FFs compared with MFs (Supplemental Figure 3) and highest in FFs when compared with IMFs present in dataset 1 (Figure 6 and Figure 7). While *FYF* expression generally decreases over time in other plants such as orchids or Arabidopsis (Chen *et al*., 2011, 2021), their expression patterns have not been explored in relation to sex expression in those species. These present expression patterns suggest that *FYF* may vary with sexual plasticity in dioecious cannabis, but further investigation is needed to determine causality. It is also possible that the discrepancy in *CsFYF*s levels between FF and other tissues is influenced by the differing lifespans of male and female cannabis flowers. Female cannabis flowers, which have a longer lifespan, may maintain higher levels of *CsFYF* to suppress senescence caused by ethylene. Exploring endogenous ethylene levels and studying the expression of *CsFYF* genes throughout flower development could provide more insights into these observed expression patterns.

Among the signaling genes analysed, *CsERS1, CsEIL1* and *CsEIL3* exhibited KC expression (Figure 5). Both IMF and FF displayed similar, lower levels of *CsERS1* expression compared to MFs, indicating a potential association between high *CsERS1* expression and male plants in *C. sativa*. However, it is worth noting that these observations contrast with a previous study by Yamasaki *et al*. (2000), which reported higher transcript levels of ethylene receptors *ETR1*, *ETR2*, and *ERS* in gynoecious cucumbers compared to monoecious cucumbers, indicating a possible connection between receptor expression and a preference for female flower production. Ethylene receptors have been the subject of extensive study in Arabidopsis, which has five isoforms of ethylene receptors. These are divided into two subfamilies, with *ERS1* and *ETR1* belongs to the first subfamily and *ERS2*, *ETR2* and *EIN4* belonging to the second subfamily (Binder, 2020). Presently no orthologs of *AtEIN4* or *AtERS2* were identified in cannabis (Figure 3). *ETR1* from subfamily 1 has largely been the focus of existing studies on ethylene perception and plant sex determination, with one study finding that *ETR1* alone could mediate silver induced inhibitions of ethylene response (McDaniel & Binder, 2012) while another study in *Cucurbita pepo* found that *ETR1* was responsible control for sex determination (García *et al*., 2020). Liu *et al*. (2012) also demonstrated that *etr1-1^(C65Y)^*Arabidopsis mutants had reduced ethylene responses independently of other receptors, highlighting the significant role of *ETR1* in ethylene signaling. However, in both datasets analysed *CsETR1* was not differentially expressed in any of the flower types. In contrast, *ERS1* which exhibited KC expression in cannabis, has been described to have a modest role as an ethylene receptor in other plants species. Studies suggest that *ERS1* may act in coordination with other ethylene receptors to affect signaling (Liu & Wen, 2012) and that its response to silver ions is much smaller than *ETR1* (McDaniel & Binder, 2012).

Ethylene insensitive-3 (*EIN3*)/Ethylene insensitive-3-like (*EIL*) family is a small family of transcription factors involved in ethylene signaling (Binder, 2020). We found that *CsEIL1* and *CsEIL3* were differentially expressed between flower types. *EILs* have been shown to play a role in physiological processes which rely on ethylene, including responses to environmental stresses (Liu *et al*., 2019), sex determination (Li *et al*., 2021) and floral development (Zhu *et al*., 2022). *EIN3/EIL1* transcription factors have been implicated with increasing the ratio of female flowers in monoecious pumpkin in response to ethephon application on seedlings (Li *et al*., 2021) and EIN3/EIL1 expression triggered by ethylene causes developmental defects and pollen sterility in anthers of *Arabidopsis.* We found higher levels of *EIL1* in XX flowers than in those of XY plants, and this raises the question as to whether elevated levels of *EIL1* detected in IMFs may affect pollen viability of induced males. In a study which overexpressed mulberry *EIL3* in Arabidopsis, the upregulation of *EIL3* also coincided with increases in ethylene biosynthesis; however, *EIL1* and *EIL2* did not follow *EIL3* expression under drought and salt stress, suggesting that expression within this family of transcription factors can vary (Liu *et al*., 2019). These findings are similar to ours, where *CsEIL1* showed clear KC expression, with highest levels of expression in FFs and IMFs, whereas *CsEIL3* showed slightly more elevated expression in MFs compared to FFs (Figure 5).

Our results highlight the differential expression of *CsERS1* and *CsEILs* between male and female flowers, supporting their involvement in sex determination mechanisms. Future studies should explore the functional significance of *ERS1* and *EILs* in sex determination pathways and investigate the cooperative interactions between *ERS1* and other ethylene receptors and downstream signaling to transcription factors in these processes. Moreover, further investigations are warranted to elucidate the specific mechanisms underlying the differential responses to silver ions among receptor subfamilies and their implications for ethylene perception.

### Genes linked to sexual plasticity found throughout the genome

*C. sativa* contains a pair of heteromorphic sex chromosomes. However, treatment of male plants with ethylene and female plants with the ethylene inhibitor STS has been reported to cause plants to produce flowers opposite than that of the sex chromosome karyotype (Green, 2005; Lubell & Brand, 2018; Moon *et al*., 2020a). The observed sexual plasticity in the species suggests that cannabis sex is not solely determined by the sex chromosomes, but rather by a combination of genetic and environmental factors (GSD+ESD), as has been suggested in the study of mammalian sexual plasticity (Lambert *et al*., 2019). There are few studies in cannabis which explore the link between sex chromosomes and sex-expression, and to the authors’ knowledge, no studies which explicitly explore molecular mechanisms linking cannabis sexual plasticity and ethylene. In a study comparing the transcriptomes of male and female flowers, Prentout *et al*. (2020) reported that 35% of sex-linked genes mapped to autosomes while 65% mapped to the sex chromosomes of *C. sativa*. The distribution of their reported sex-linked genes in the chromosome is in stark contrast to the distribution of CsERGs that we identified with only 26.2% of CsERGs mapping to the sex chromosomes. When considering only CsERGs which are differentially expressed, 20% (3/15) are located on the sex chromosomes, while the remainder are located on the autosomes (Figure 4). The ubiquity of ERGs in the autosomes is to be expected, as they play important roles in general plant growth with ethylene regulating many plant growth responses. Prentout *et al*. (2020) suggested that their observation of autosomal mapping for sex-linked genes might have been the result of false positives or incorrect placement of genes in the genome assembly of *C. sativa*. However, their assumption appears to overlook the possibility that many genes involved in plant sex might also be crucial for the production of essential growth regulators or metabolites, which are required in both genetically male and female *C. sativa* and whose loci are predominantly found on autosomes. In contrast, Adal *et al*. (2021) proposes an X:autosome dosage mechanism as the basis for sex determination, where male flowers might possess a limited number of unique Y-chromosome genes influencing their sex. We propose that the regulation of cannabis sexual plasticity is mediated by the differential expression of CsERGs, encompassing both FOC and uERG expression, on both autosomes and sex chromosomes of cannabis. The presence of CsERGs on autosomes aligns with ethylene’s involvement in general plant metabolism, while their presence on sex chromosomes suggests some specific roles in sexual development. By considering the comprehensive distribution of CsERGs across the genome, we highlight the importance of both autosomal and sex chromosome-associated genes in the regulation of cannabis sexual plasticity.

### Limitations and future directions

Based on previous observations suggesting a link between ethylene and sexual plasticity/determinism, the present study constitutes a preliminary analysis to identify putative candidate genes involved in the sexual expression of cannabis. However, in the case of this exploratory study, there are some limitations that we would like to acknowledge. Only two transcriptomic datasets from floral tissues were available that contained multiple flower sexes (Prentout *et al*., 2020; Adal *et al*., 2021). In an effort to begin probing the effect of CsERGs during chemically induced sexual plasticity, it was important that some of the transcriptomes include IMFs, which were only available in one of the two datasets used (Adal *et al*., 2021). Additionally, no existing transcriptomes were available that contained IFFs, which limited our ability to test whether there is an inverse relationship in the mechanisms for sexual plasticity between MFs and IFFs. Future studies on sexual plasticity in cannabis should include IFFs in order to provide a more complete picture of the mechanisms underlying sexual plasticity in the species. Despite these limitations, the differential expression findings between MF and FF in both datasets was used to find a set of differentially expressed CsERGs (Supplemental Figure 1 and Supplemental Figure 3), which are strong candidate CsERGs with a role in sex determination.

One limitation of the study is the use of datasets collected at a single timepoint, specifically from flowers at a later stage of development. This approach may result in the potential omission of differentially expressed CsERG candidates that are not active during that specific timepoint, particularly those involved in early floral sex determination or sex differentiation. Furthermore, it is important to acknowledge that some of the identified candidates may exhibit differential expression at the selected timepoint for reasons unrelated to the intended analysis, which aims to establish their role in sexuality. The datasets employed in this study were generated under different experimental conditions, including variations in plant growth conditions, sampling stages of flower development, and sampling locations on the plants. It is crucial to note that the datasets were derived from distinct genotypes of *C. sativa*: one drug-type (cv. MS-17-338; Adal *et al*., 2021) and one hemp-type cannabis (cv. Zenitsa; Prentout *et al*., 2020). As cannabis is known for its high genetic variability (Lapierre *et al*., 2023), studies based on a single genotype lack the robustness necessary for generalization (Page *et al*., 2021; Monthony *et al*., 2021b). Therefore, future studies on *C. sativa* should aim to include a broader range of cannabis genotypes, particularly those exploring the role of ethylene in plasticity and sex determination. Notably, despite these limitations, the transcriptomes from the two different cannabis populations facilitated the validation of 34 out of the 43 predicted CsERGs listed in Table 1. This validation adds robustness to the gene models proposed in this study for the Yang Cycle, ethylene biosynthesis, and signaling pathways (Figure 2 and Figure 3). It is important to emphasize that the lack of expression of certain predicted CsERGs in both datasets does not necessarily imply that they are non-functional CsERGs. As previously discussed, the inclusion of transcriptomes covering a wider timeline, including early floral induction stages, will contribute to the comprehensive validation of these predicted models.

There are currently a limited number of biotechnological tools (e.g., *in vitro* culture, CRISPR, VIGS, agrobacterium, etc.) which are optimized in order to functionally validate CsERGs in cannabis (Hesami *et al*., 2021) and many of these tools have been hampered by the lack of a reliable *in vitro* regeneration protocol for the species (Monthony *et al*., 2021b). Although such functional validation is beyond the scope of the current study, it represents an important next step in probing the role of ethylene in cannabis sexual plasticity. Recently progress has been made in establishing methods for gene silencing in cannabis, through the employment of the *Cotton leaf crumple virus* (*CLCrV*) in a virus-induced gene silencing protocol reported by Schachtsiek *et al*. (2019) and through new methods using agrobacterium (Sorokin *et al*., 2020; Galán-Ávila *et al*., 2021). However due to their recent nature, these methods have yet to be replicated independently. Given the genotypic variability of cannabis and the role it plays in the replicability of *in vitro* methods (Monthony *et al*., 2021b), it remains to be seem how effectively these methods are adapted to multiple genotypes across different research settings.

## Conclusion

This study represents the first step in exploring sexual plasticity in the dioecious plant *C. sativa* and highlights the importance of the Yang cycle, ethylene biosynthesis and signaling pathways in sex determination. Here, we present a framework consisting of three distinct gene expression patterns (KC, FOC and uERG) which can be used to categorize the role of CsERGs in sexual plasticity in dioecious plants. These findings are based on our analysis of transcriptomes of male, female and induced male flowers obtained from two different *Cannabis sativa* genotypes produced under two different experimental conditions. From the ortholog analysis undertaken in this study and with support from transcriptomic data, we propose that the canonical Yang Cycle and ethylene biosynthesis and signaling pathways are conserved in *C. sativa* and we identified seven genes belonging to two the FOC and uERG classes which we hypothesize play a role in ethylene-induced sexual plasticity in cannabis. This work lays the groundwork for more detailed study of the role of CsERGs in sexual plasticity in multiple genotypes of cannabis. Given the commercial importance of cannabis, as well as a number of other dioecious plants such as kiwi, papaya and persimmon, understanding the underlying factors of sexual plasticity and its link with ethylene are important to future efforts in breeding and crop improvement.

## Acknowledgments

The authors gratefully acknowledge the support of the Natural Sciences and Engineering Research Council (NSERC) of Canada. ASM has also been supported by a NSERC Canada Vanier Graduate Scholarship. The authors would also like to thank Eric Normandeau for his support with script writing.

## Author Contributions

ASM, MR, and DT planned and designed the research. ASM and MR performed the investigation and contributed to data curation. ASM and MR conducted formal analysis. ASM and DT acquired funding for the project. ASM, MR, and DT contributed to the methodology. ASM and DT handled project administration. DT provided resources. DT supervised the research. ASM visualized the results. ASM and DT wrote the original draft of the manuscript. ASM, MR, and DT contributed to reviewing and editing the manuscript.

## Data Availability Statement

All scripts and raw data used in this study are available from The Open Science Framework (OSF) under the DOI: 10.17605/OSF.IO/7G9VR and at https://osf.io/7g9vr/?view_only=b633db1f598a4d31a975c8293eb2a299. Transcriptomes were previously published on the NCBI under the accession numbers: PRJNA669389 (dataset 1) and PRJNA549804 (dataset 2).

