## Supplemental Tables and Figures v2 for "Exploring Ethylene-Related Genes in *Cannabis sativa*: Implications for Sexual Plasticity"

### Supplementary Tables

**Supplementary Table 1** Summary of RNA seq data used for differential expression analysis of CsERGs. RNA was obtained from three flower types (male flowers, MF; induced male flowers, IMF; and female flowers, FF). SRA accession numbers and biological replicates are listed. SRA accession numbers beginning with SRR are from dataset 1 and SRX from dataset 2.

| Flower type | SRA accession numbers | Biological replicates | Dataset |
|---|---|---|---|
| MF | SRR12831871, 72 | 2 | 1 |
| | SRX6096794, 795, 801, 802, 804 | 5 | 2 |
| IMF | SRR12831863*, 64, 65, 75 | 4 | 1 |
| FF | SRR12831869, 70, 76 | 3 | 1 |
| | SRX6096793, 796, 799, 800, 803 | 5 | 2 |

*Indicates samples were excluded from DE analysis due to low quality mapped reads GC-content distribution.

**Supplementary Table 2** A total of 47 Arabidopsis derived ethylene-related genes (ERG) were identified and used for homology analysis to search for CsERGs in the *C. sativa* Cs10 reference genome.

| Arabidopsis ERG | Locus ID | Pathway | Source |
|---|---|---|---|
| ACO1 | AT2G19590 | Ethylene Biosynthesis | (Moon *et al.*, 2020b) |
| ACO2 | AT1G62380 | Ethylene Biosynthesis | (Raz & Ecker, 1999) |
| ACO3 | AT1G12010 | Ethylene Biosynthesis | (Cheng *et al.*, 2009) |
| ACO4 | AT1G05010 | Ethylene Biosynthesis | (Moon *et al.*, 2020b) |
| ACO5 | AT1G77330 | Ethylene Biosynthesis | (Brenner *et al.*, 2005) |
| ACS2 | AT1G01480 | Ethylene Biosynthesis | (Yamagami *et al.*, 2003) |
| ACS4 | AT2G22810 | Ethylene Biosynthesis | (Yamagami *et al.*, 2003) |
| ACS5 | AT5G65800 | Ethylene Biosynthesis | (Yamagami *et al.*, 2003) |
| ACS6 | AT4G11280 | Ethylene Biosynthesis | (Yamagami *et al.*, 2003) |
| ACS7 | AT4G26200 | Ethylene Biosynthesis | (Yamagami *et al.*, 2003) |
| ACS8 | AT4G37770 | Ethylene Biosynthesis | (Yamagami *et al.*, 2003) |
| ACS9 | AT3G49700 | Ethylene Biosynthesis | (Yamagami *et al.*, 2003) |
| ACS11 | AT4G08040 | Ethylene Biosynthesis | (Yamagami *et al.*, 2003) |
| ARD1 | AT4G14716 | Yang Cycle | (Bürstenbinder *et al.*, 2007) |
| ARD2 | AT4G14710 | Yang Cycle | (Bürstenbinder *et al.*, 2007) |
| ARD3 | AT2G26400 | Yang Cycle | (Bürstenbinder *et al.*, 2007) |
| ARD4 | AT5G43850 | Yang Cycle | (Bürstenbinder *et al.*, 2007) |
| ARGOS* | AT3G59900 | Ethylene Adjacent | (Shi *et al.*, 2015) |

| | | | |
|---|---|---|---|
| CTR1 | AT5G03730 | Ethylene Signaling | (Kieber *et al.*, 1993) |
| DEP1 | AT5G53850 | Yang Cycle | (Pommerrenig *et al.*, 2011) |
| EBF1 | AT2G25490 | Ethylene Signaling | (Guo & Ecker, 2003) |
| EBF2 | AT5G25350 | Ethylene Signaling | (Guo & Ecker, 2003) |
| EIL1 | AT2G27050 | Ethylene Signaling | (Chao *et al.*, 1997) |
| EIL2* | AT5G21120 | Ethylene Signaling | (Chao *et al.*, 1997) |
| EIL3 | AT1G73730 | Ethylene Signaling | (Chao *et al.*, 1997) |
| EIN2 | AT5G03280 | Ethylene Signaling | (Guzman & Ecker, 1990) |
| EIN3 | AT3G20770 | Ethylene Signaling | (Roman *et al.*, 1995) |
| EIN4 | AT3G04580 | Ethylene Signaling | (Roman *et al.*, 1995) |
| ERF1 | AT3G23240 | Ethylene Signaling | (Solano *et al.*, 1998) |
| ERS1 | AT2G40940 | Ethylene Signaling | (Sakai *et al.*, 1998) |
| ERS2 | AT1G04310 | Ethylene Signaling | (Sakai *et al.*, 1998) |
| ETP1 | AT3G18980 | Ethylene Signaling | (Qiao *et al.*, 2009) |
| ETP2 | AT3G18910 | Ethylene Signaling | (Qiao *et al.*, 2009) |
| ETR1 | AT1G66340 | Ethylene Signaling | (Bleecker *et al.*, 1988) |
| ETR2 | AT3G23150 | Ethylene Signaling | (Sakai *et al.*, 1998) |
| FYF | AT5G62165 | Ethylene Adjacent | (Chen *et al.*, 2011) |
| MTI1 | AT2G05830 | Yang Cycle | (Pommerrenig *et al.*, 2011) |
| MTK1 | AT1G49820 | Yang Cycle | (Pommerrenig *et al.*, 2011) |
| MTN1 | AT4G38800 | Yang Cycle | (Pommerrenig *et al.*, 2011) |
| MTN2 | AT4G34840 | Yang Cycle | (Pommerrenig *et al.*, 2011) |
| ORE1 | AT5G39610 | Ethylene Adjacent | (Qiu *et al.*, 2015) |
| RAN1 | AT5G44790 | Ethylene Signaling | (Hirayama *et al.*, 1999) |
| RTE1 | AT2G26070 | Ethylene Signaling | (Resnick *et al.*, 2006) |
| MAT1 (SAMS1) | AT1G02500 | Yang Cycle | (Peleman *et al.*, 1989a) |
| MAT2 (SAMS2) | AT4G01850 | Yang Cycle | (Peleman *et al.*, 1989b). |
| MAT3 (SAMS3) | AT2G36880 | Yang Cycle | (Chen *et al.*, 2016b) |
| MAT4 (SAMS4) | AT3G17390 | Yang Cycle | (Meng *et al.*, 2018) |

* denotes AtERGs for which no CsERG orthologs were found during the ortholog analysis.

**Supplementary Table 3** Results of an InterProScan of the CsERGs. Protein family is given when available, otherwise the conserved protein domains are provided, alongside the InterPro entry for each family/domain. Graphical representations of InterProScan results for each CsERG can be viewed on the OSF database for this article.

| CsERG Locus | CsERG Name | Protein Family/ Conserved Domains | InterPro Entry |
|---|---|---|---|
| LOC115704991 | CsACO1 | Oxoglutarate/iron-dependent dioxygenase* | IPR005123 |

| | | | |
|---|---|---|---|
| | | Non-haem dioxygenase N-terminal domain* | IPR026992 |
| | | Isopenicillin N synthase-like, Fe(2+) 2OG dioxygenase domain* | IPR044861 |
| LOC115702109 | CsACO1-LIKE1 | Oxoglutarate/iron-dependent dioxygenase* | IPR005123 |
| | | Isopenicillin N synthase-like, Fe(2+) 2OG dioxygenase domain* | IPR044861 |
| | | Non-haem dioxygenase N-terminal domain* | IPR026992 |
| LOC115707397 | CsACO2 | Non-haem dioxygenase N-terminal domain* | IPR026992 |
| | | Isopenicillin N synthase-like, Fe(2+) 2OG dioxygenase domain* | IPR044861 |
| | | Oxoglutarate/iron-dependent dioxygenase* | IPR005123 |
| LOC115699412 | CsACO3 | Oxoglutarate/iron-dependent dioxygenase* | IPR005123 |
| | | Non-haem dioxygenase N-terminal domain* | IPR026992 |
| | | Isopenicillin N synthase-like, Fe(2+) 2OG dioxygenase domain* | IPR044861 |
| LOC115699408 | CsACO5 | Isopenicillin N synthase-like, Fe(2+) 2OG dioxygenase domain* | IPR044861 |
| | | Oxoglutarate/iron-dependent dioxygenase* | IPR005123 |
| | | Non-haem dioxygenase N-terminal domain* | IPR026992 |
| LOC115704849 | CsACS1 | Aminotransferase, class I/class II* | IPR004839 |
| LOC115717044 | CsACS2 | Aminotransferase, class I/class II* | IPR004839 |
| LOC115712396 | CsACS3 | Aminotransferase, class I/class II* | IPR004839 |

| | | | |
|---|---|---|---|
| LOC115704963 | CsACS7-LIKE | Aminotransferase, class I/class II* | IPR004839 |
| LOC115696400 | CsACS9 | Aminotransferase, class I/class II* | IPR004839 |
| LOC115716540 | CsACS9-LIKE | Aminotransferase, class I/classII* | IPR004839 |
| LOC115721731 | CsARD1 | Acireductone dioxygenase ARD family* | IPR004313 |
| | | Acireductone dioxygenase, eukaryotes* | IPR027496 |
| LOC115722194 | CsARD2 | Acireductone dioxygenase ARD family* | IPR004313 |
| | | Acireductone dioxygenase, eukaryotes* | IPR027496 |
| LOC115697744 | CsCTR1 | Protein kinase domain* | IPR000719 |
| | | Serine-threonine/tyrosine-protein kinase, catalytic domain* | IPR001245 |
| LOC115706236 | CsCTR1-LIKE | Protein kinase domain* | IPR000719 |
| | | Serine-threonine/tyrosine-protein kinase, catalytic domain* | IPR001245 |
| LOC115706293 | CsDEP1 | Enolase-phosphatase E1* | IPR023943 |
| | | Methylthioribulose-1-phosphate dehydratase, eukaryotes* | IPR027514 |
| | | Methylthioribulose-1-phosphate dehydratase* | IPR017714 |
| | | Probable bifunctional methylthioribulose-1-phosphate dehydratase/enolase-phosphatase E1* | IPR027505 |
| LOC115720532 | CsEBF1 | F-box domain* | IPR001810 |
| LOC115724585 | CsEBF2 | F-box domain* | IPR001810 |
| LOC115711236 | CsEIL1 | Ethylene insensitive 3* | IPR006957 |
| LOC115719373 | CsEIL3 | Ethylene insensitive 3* | IPR006957 |
| LOC115709621 | CsEIN2 | NRAMP family* | IPR001046 |
| | | Ethylene-insensitive protein 2* | IPR017187 |
| LOC115706079 | CsEIN3 | Ethylene insensitive 3* | IPR006957 |

| | | | |
|---|---|---|---|
| LOC115723050 | CsERF1 | AP2/ERF domain* | IPR001471 |
| LOC115697291 | CsERS1 | Signal transduction histidine kinase, dimerisation/phosphoacceptor domain* | IPR003661 |
| | | Signal transduction histidine kinase-related protein, C-terminal* | IPR004358 |
| | | GAF domain* | IPR003018 |
| | | Histidine kinase/HSP90-like ATPase* | IPR003594 |
| | | Histidine kinase domain* | IPR005467 |
| LOC115705474 | CsETP1 | F-box associated domain, type 1* | IPR006527 |
| | | F-box domain* | IPR001810 |
| | | F-box associated interaction domain* | IPR017451 |
| LOC115720323 | CsETP2 | F-box domain* | IPR001810 |
| | | F-box associated interaction domain* | IPR017451 |
| | | F-box associated domain, type 3 | IPR013187 |
| LOC115702329 | CsETR1 | Ethylene receptor* | IPR014525 |
| LOC115721785 | CsETR2 | GAF domain* | IPR003018 |
| | | Signal transduction response regulator, receiver domain* | IPR001789 |
| | | Signal transduction histidine kinase, dimerisation/phosphoacceptor domain* | IPR003661 |
| | | Histidine kinase domain | IPR005467 |
| LOC115700576 | CsFYF1 | MADS MEF2-like* | IPR033896 |
| | | Transcription factor, MADS-box* | IPR002100 |
| | | Transcription factor, K-box* | IPR002487 |
| LOC115706939 | CsFYF2 | Transcription factor, K-box* | IPR002487 |
| | | Transcription factor, MADS-box* | IPR002100 |
| | | MADS MEF2-like* | IPR033896 |
| LOC115716986 | CsFYF3 | Transcription factor, K-box* | IPR002487 |

| | | | |
|---|---|---|---|
| | | Transcription factor, MADS-box* | IPR002100 |
| | | MADS MEF2-like* | IPR033896 |
| LOC115717309 | CsFYF4 | Transcription factor, K-box* | IPR002487 |
| | | Transcription factor, MADS-box* | IPR002100 |
| | | MADS MEF2-like* | IPR033896 |
| LOC115711364 | CsMAT1 | S-adenosylmethionine synthetase* | IPR002133 |
| LOC115719259 | CsMAT2 | S-adenosylmethionine synthetase* | IPR002133 |
| LOC115711343 | CsMAT3 | S-adenosylmethionine synthetase* | IPR002133 |
| LOC115706479 | CsMTI1 | Methylthioribose-1-phosphate isomerase* | IPR005251 |
| | | Initiation factor 2B alpha/beta/delta* | IPR011559 |
| | | Initiation factor 2B-related* | IPR000649 |
| LOC115703014 | CsMTK1 | Methylthioribose kinase* | IPR009212 |
| LOC115702983 | CsMTK1-LIKE | Methylthioribose kinase* | IPR009212 |
| LOC115704327 | CsMTN | 5'-Methylthioadenosine/S-adenosylhomocysteine nucleosidase* | IPR044580 |
| LOC115701489 | CsORE1 | NAC domain* | IPR003441 |
| LOC115709772 | CsORE2 | NAC domain* | IPR003441 |
| LOC115698184 | CsRAN1 | P-type ATPase* | IPR001757 |
| | | P-type ATPase, subfamily IB* | IPR027256 |
| LOC115707134 | CsRTE1 | TMEM222/RTE1* | IPR008496 |

* indicates that the protein family/domain was present in the Arabidopsis ortholog of the CsERG.

**Supplementary Table 4** Distribution of the 43 CsERGs across the *C. sativa* cs10 reference genome, based on their known functions in relation to ethylene.

| Chromosome | Biosynthesis % (count) | Ethylene Adjacent % (count) | Signaling % (count) | Yang Cycle % (count) |
|---|---|---|---|---|
| Chr1 | 40 (4) | 16.6 (1) | 17.6 (3) | 20 (2) |
| Chr2 | 0 | 0 | 11.8 (2) | 10 (1) |
| Chr3 | 0 | 16.6 (1) | 5.9 (1) | 10 (1) |

| | | | | |
|---|---|---|---|---|
| Chr4 | 10 (1) | 0 | 0 | 0 |
| Chr5 | 20 (2) | 33.3 (2) | 0 | 0 |
| Chr6 | 0 | 0 | 5.9 (1) | 0 |
| Chr7 | 0 | 0 | 11.8 (2) | 0 |
| Chr8 | 10 (1) | 33.3 (2) | 0 | 0 |
| Chr9 | 0 | 0 | 11.8 (2) | 20 (2) |
| ChrX | 20 (2) | 0 | 29.4 (5) | 40 (4) |
| Unassigned Scaffold | 0 | 0 | 5.9 (1) | 0 |

### Supplementary Figures

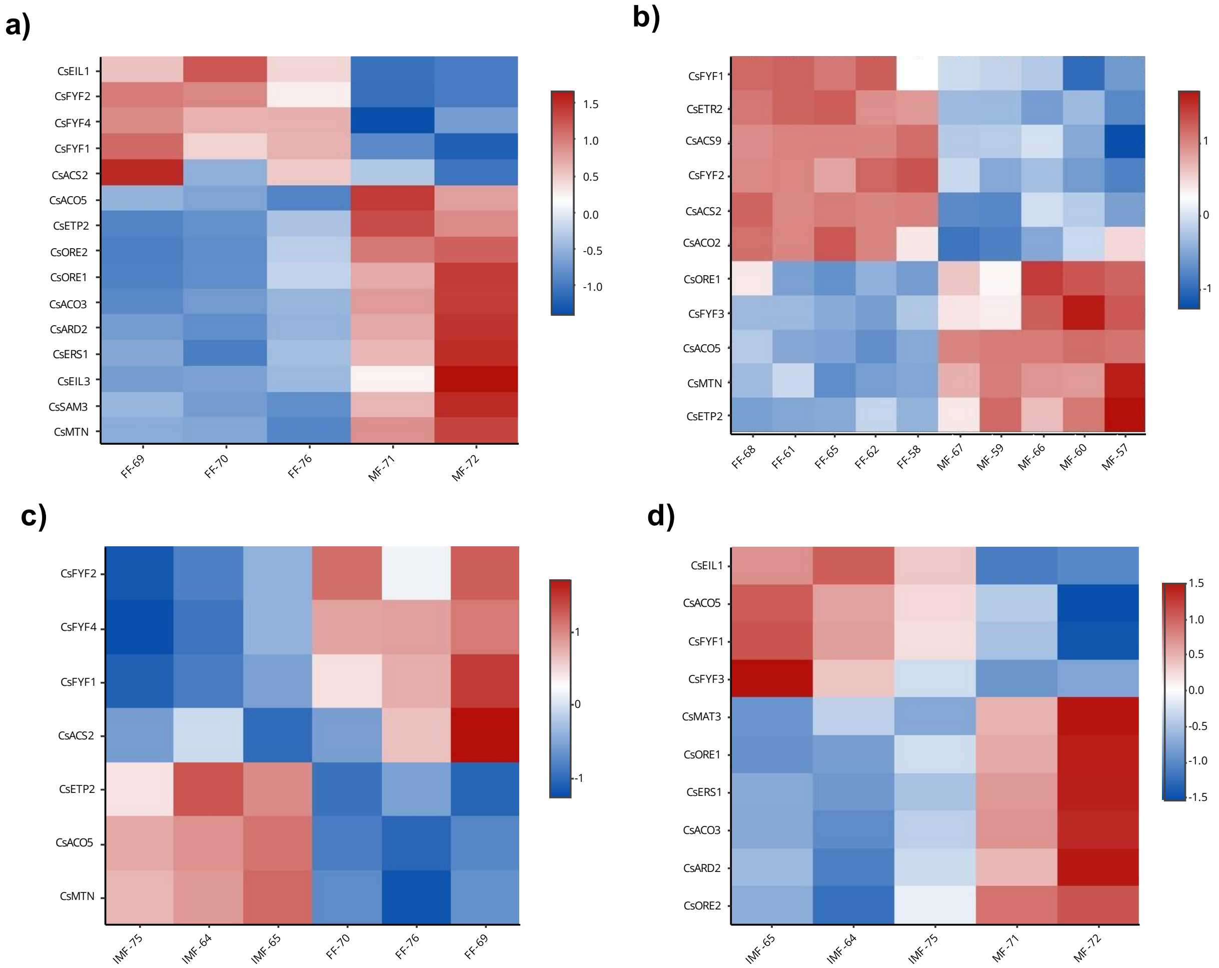


**Supplemental Figure 1** Heatmaps of differentially expressed CsERGs (identified by CsERG name; **Table 1**) for dataset 1 (a, c and d) and dataset 2 (b). X-axis labels are the flower type, followed by the last two digits of the SRA accession numbers, which can be found in Supplementary Table 1. Y-axis contains the CsERG names. Heatmap scale is in log of the fold change in expression for the pairwise comparison of the flower type listed on the x-axis: a) MF vs. FF comparison in dataset 1; b) MF vs. FF comparison in dataset 2; c) IMF vs. FF in dataset 1; d) IMF vs. MF in dataset 1.

44

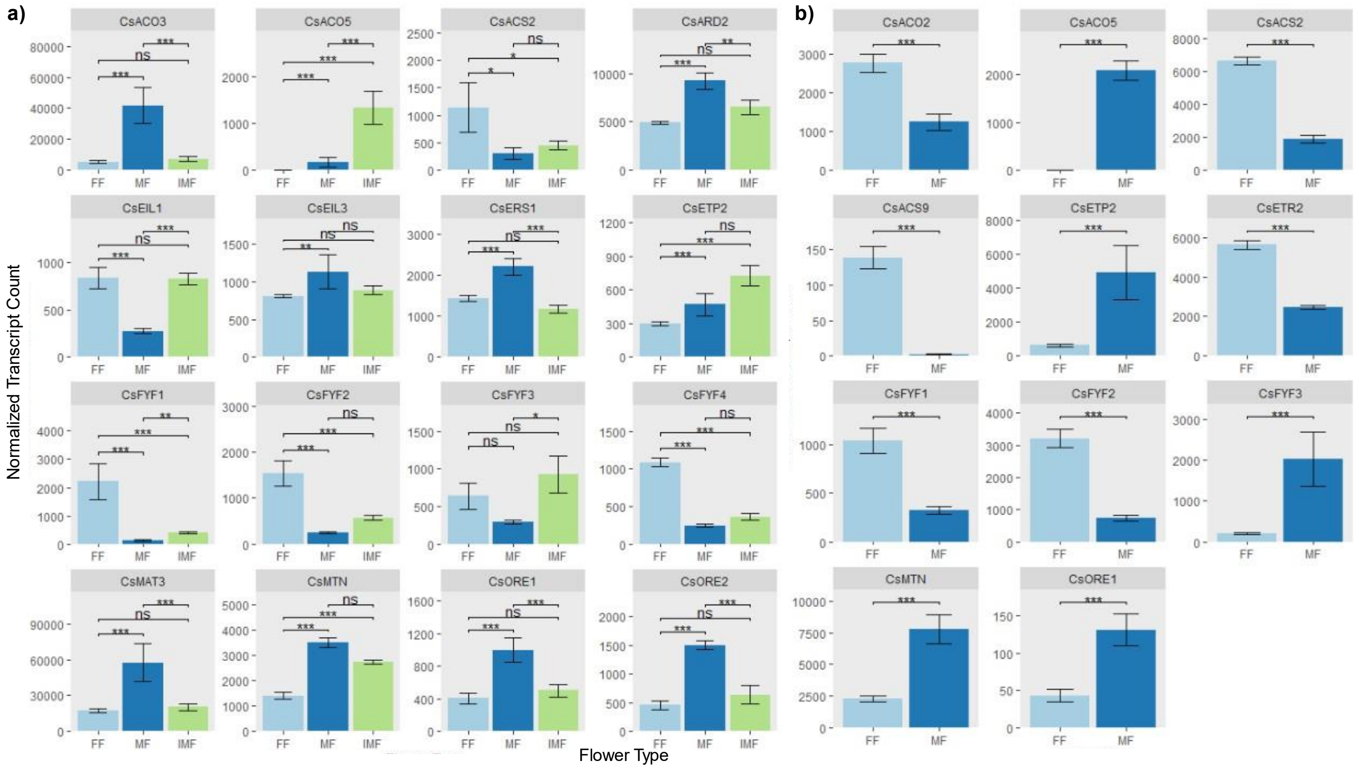


45

**Supplemental Figure 2** CsERG gene expression levels for CsERGs where differential expression was detected in dataset 1 (a) and dataset 2 (b). *C. sativa* gene names are indicated on the top of all panels, x-axes show floral treatments (female flower, FF; male flower, MF; induced male flower, IMF), y-axes show normalized transcript counts. Pairwise comparisons of CsERGs were considered significantly different if they had an adjusted p-value of $<0.05$ and a $|\log2foldchange| > 1$. Significant pairwise comparisons are denoted by asterisks: ns = not significant; * for $p \leq 0.05$; ** for $p \leq 0.01$; *** for $\leq 0.001$ as determined by the Wald test.

46

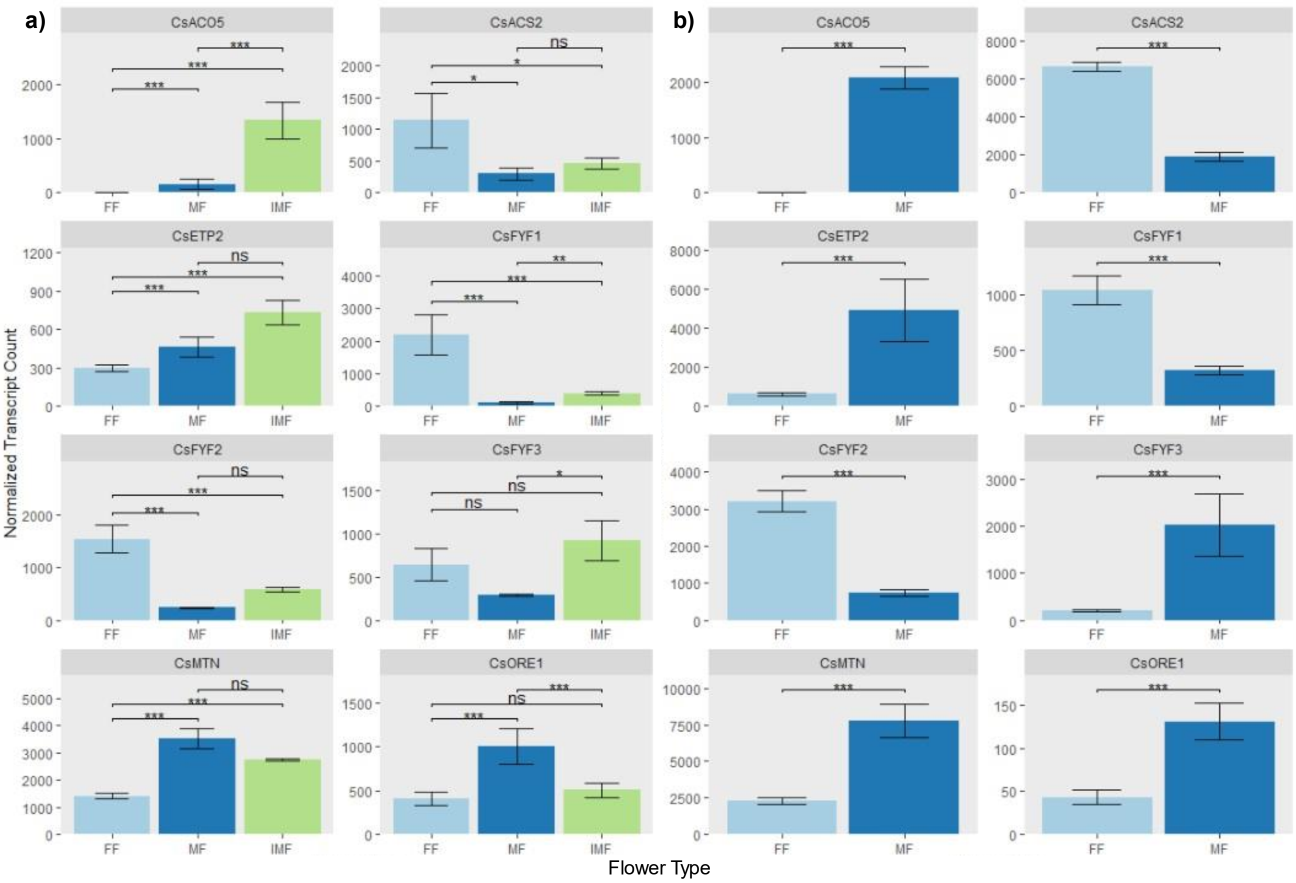


**Supplemental Figure 3** CsERGs which showed similar patterns of differential expression between FF and MF flower types in dataset 1 and dataset 2. Significant pairwise comparisons are denoted by asterisks: ns = not significant; * for $p \leq 0.05$; ** for $p \leq 0.01$; *** for $\leq 0.001$ as determined by the Wald test.

### BLASTP identification of orthologs

All files that were used in the identification of orthologs using BLASTP and that were not publicly accessible have been uploaded to a database on the Open Science Framework at: https://osf.io/7g9vr/?view_only=b633db1f598a4d31a975c8293eb2a299. A local BLASTP was performed using the cs10 cannabis reference proteome as the database (RefSeq accession: GCF_900626175.2) obtained from the NCBI. The query list of AtERGs proteins sequences was obtained from the gene list in **Supplementary Table *2*** (also available as a gene collection on NCBI at: https://www.ncbi.nlm.nih.gov/sites/myncbi/adrian.monthony.1/collections/61572191/public/) and the corresponding protein sequences were obtained by entering the *A. thaliana* gene IDs into the data table, opening transcript view and downloading the FASTA protein sequences using the

NCBI Datasets page (https://www.ncbi.nlm.nih.gov/datasets/tables/genes/). Files containing the *A. thaliana* gene and protein sequences can be found on the OSF database for this project. Following the BLASTP duplicate BLASPT results interrupted by introns or non-coding regions were merged using a custom python script which can also be found on the OSF page for this study. The results from the BLASTP were formatted using the -outfmt "6 qseqid sseqid pident length mismatch gapopen qstart qend sstart send evalue bitscore stitle" condition. The formatted results were subsequently split into individual .tsv files, one for each *A. thaliana* protein sequence query, so that for each query putative orthologs in the cs10 proteome could be identified. Creation of individual files for each Arabidopsis protein BLASTP result was done using the Genebygene_sample_script.sh available on the OSF. Each result was manually analyzed for orthologs by plotting three, 2-way comparisons between sequence length, similarity and bit score using RStudio (script available on OSF). Orthologous protein sequences in the cs10 genome were identified when they showed high similarity, sequence length and bit score in the 2-way comparison (Supplemental Figure 4a). When no high-scoring putative results were found for a given Arabidopsis protein query, it was assumed that no orthologs existed (Supplemental Figure 4b). In the case of ERGs for which no high-scoring matches were found using BLASTP, a second manual examination found mid-scoring results which were used as putative orthologs. This was done in the case of *AtETP1/2*, for which no *C. sativa* orthologs were found following an initial OrthoFinder or BLASTP analysis. One possible explanation is that the numerous highly conserved F-BOX proteins found in the protein family may have made high scoring and high similarities between *ETP1/2* protein sequences in *A. thaliana* and *C. sativa* difficult to distinguish during ortholog analysis, leading to no obvious orthologs. A second manual examination of the BLASTP results for *AtETP1* and *AtETP2* identified for each a single result that stood out from the other results because of long sequence length vs. bit score profiles, despite lower percentage similarities than other for other CsERGs identified using the BLASTP in this study (see Supplemental Figure 4c).

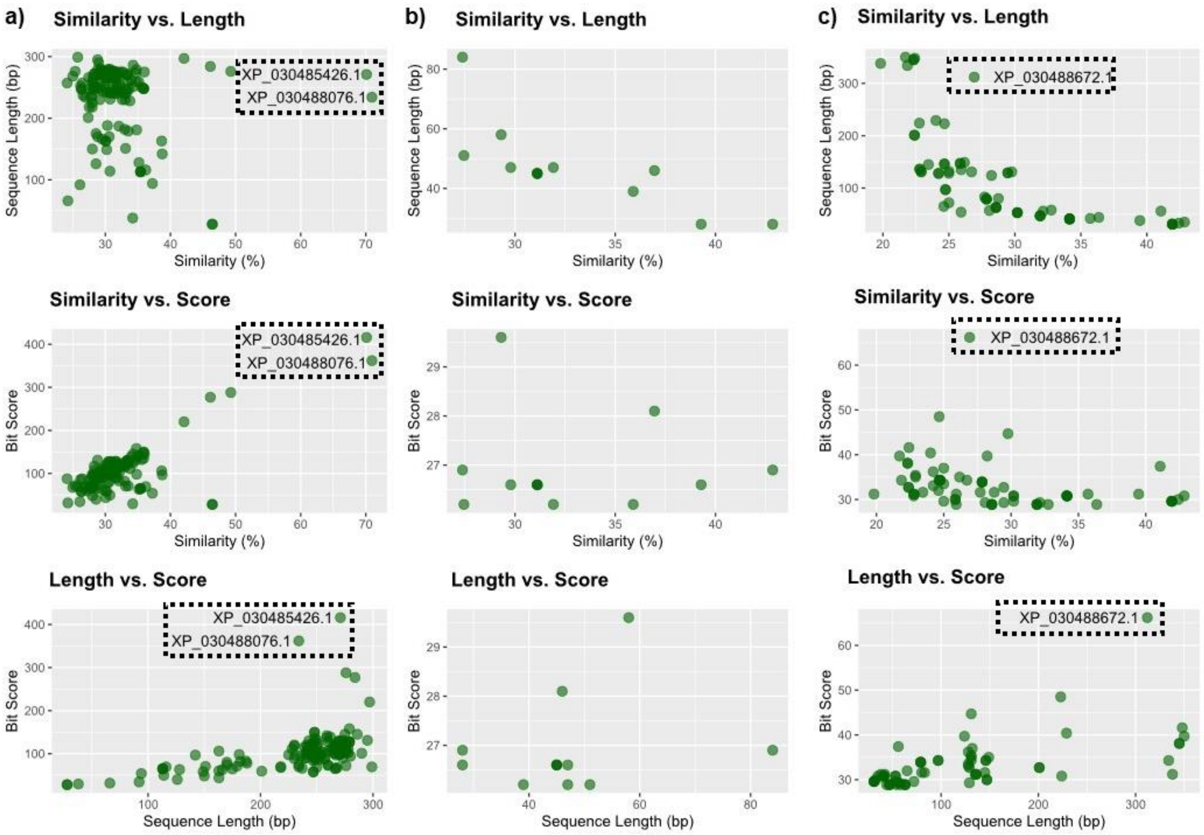


**Supplemental Figure 4** This figure shows examples of putative orthologs identification, for high-scoring CsERG proteins (a), proteins for which no orthologs were identified (b) and proteins for which a second manual analysis of the data was required in order to identify a putative ortholog (c). Putative orthologs are identified in the dashed box based on results from three, two-way comparisons of BLASTP results: similarity (%), length (bp) and bit score. a) NP_001324863 (AT2G19590; *A. thaliana* ACC oxidase, b) shows an Arabidopsis protein NP_191549.2 (AT3G59900; *A. thaliana* ARGOS) for which no cs10 orthologs were identified using BLASTP and c) shows mid-scoring putative ortholog of *A. thaliana* ETP1(AT3G18980; NP_001078185.1) which was identified upon a second review of the BLASTP data.
